# Senescent CD8 T Effector Memory Cells are Functionally Impaired, Enriched in Aging and Disease, and a Barrier to Immunotherapy

**DOI:** 10.64898/2025.12.16.694716

**Authors:** Paolo S. Turano, Elizabeth Akbulut, Nica Mariz Aquino, Luis Garza-Martínez, Sukhwinder Singh, George S. Yap, Patricia Fitzgerald-Bocarsly, Ricardo Iván Martínez-Zamudio, Utz Herbig

## Abstract

Senescent cells play important roles in various biological processes that promote fitness and health, however, their timely elimination by immune cells is critical to maintain tissue homeostasis and prevent disease. Despite this, senescent cells progressively accumulate systemically with age, suggesting that certain immune cells also become senescent and dysfunctional during aging. Supporting this, we previously demonstrated that CD8 T cells, immune cells capable of targeting senescent cells, increasingly develop characteristics of senescence with advancing age in humans. Here, we further characterized the senescence state of human SA-βGal-expressing CD8 T effector cells, their functional capabilities, and their involvement in aging and disease. Single-cell RNA sequencing revealed that SA-βGal-expressing CD8 T cells with unique transcriptional signatures develop in all stages of T cell differentiation, including in effector memory (EM) T cells. SA-βGal-expressing CD8 T_EM_ cells expressed various classical markers of senescence and were significantly impaired in their ability to proliferate, produce cytokines, and eliminate senescent human stromal cells, compared to CD8 T_EM_ cells with low SA-βGal activity. Gene signatures of senescent SA-βGal-expressing CD8 T_EM_ cells were enriched in CD8 T cells from older human donors, patients with age-related disorders, cancer, and smokers. Furthermore, our results demonstrate that T cell senescence is distinct from and dominant over T cell exhaustion, limiting the response of CD8 T_EM_ cells to immunotherapy. Collectively, our study demonstrates that the senescence state impairs the functions of CD8 T_EM_ cells and reveals the involvement of senescent and dysfunctional CD8 T_EM_ cells in aging, disease, exposure to toxins, and responses to immunotherapy.

## Introduction

Cellular senescence is a state in which cells can no longer divide, exhibit altered or impaired functions, and release a mix of bioactive molecules that can influence the surrounding extracellular environment^1^. Although senescent cells play important roles in diverse biological processes that promote fitness and health, their timely elimination by immune cells is crucial for maintaining tissue homeostasis and preventing disease^2^. As we age, however, immune-mediated clearance of senescent cells becomes increasingly less efficient, as evidenced by the observed age-associated and systemic increase of senescent cells. This age-associated accumulation of senescent cells has a profound negative impact on health, as it promotes the development of various age-related disorders and diseases, including cancer^3^. Why our immune system increasingly fails to eliminate senescent cells with age remains unclear.

A decline in immune function with age, termed immunosenescence, has been attributed to several factors, including age-associated impairment in thymopoiesis, a reduction in the ratio of naïve to memory cells, and a progressive increase in chronic, low-grade inflammation^4^. As various immune cells increasingly develop features of cellular senescence with age, it is possible that immune cell senescence also contributes to immunosenescence and systemic accumulation of senescent cells. This is supported by studies demonstrating that mice with impaired T cell function^5,6^, or with autologously transplanted aged or senescent splenocytes^7^, develop significantly more non-lymphoid senescent cells systemically during aging, resulting in an increased development of various age-related pathologies and reduced lifespans.

The greatest accumulation of senescent immune cells in humans has been reported in the CD8 T cell compartment^8^. CD8 T cells are cytotoxic immune cells that can directly migrate towards and target aberrant and senescent non-lymphoid cells^9–13^ by releasing perforin and granzymes, thereby triggering cell death. In human peripheral blood, CD8 T cells exist in various differentiation stages, including naïve, antigen-inexperienced cells (T_N_), as well as in central memory (T_CM_) and effector memory (T_EM_) states, which are long-lived, antigen-experienced cells that can mount a rapid cytotoxic response upon re-encountering their specific target antigens. A fourth CD8 T cell subset, called T_EMRA_ (effector memory re-expressing CD45RA), develops in response to repeated antigen exposure and consists of terminally differentiated cells with low proliferative capacity but high effector function, resembling a natural killer (NK) cell-like phenotype^14,15^. Although it has been argued that CD8 T_EMRA_ cells are in a state that most closely resembles cellular senescence^16^, subpopulations of T_EMRA_ cells retain the ability to proliferate^17^ and lack classical features of cellular senescence^8^, suggesting that this cell state is only partially comprised of senescent cells^18^.

Another dysfunctional T cell state, known as T cell exhaustion, develops in response to persistent antigenic stimulation and is characterized by the expression of inhibitory receptors, such as PD-1^19,20^. T cell exhaustion is associated not only with aging^12^ but also with the failure of T cells to target and eliminate cancer cells^19^. Blocking the interaction of these inhibitory receptors with their respective ligands using receptor-specific antibodies, a process called immune checkpoint blockade (ICB), allows exhausted T cells to overcome their functional impairment and kill target cells. While these ICB therapies are used in the clinic to treat various cancers, they show mixed results, as many patients do not respond to ICB for reasons that are still incompletely understood^19^.

Although senescence can be triggered by various cell-intrinsic and extrinsic signals, a primary cause of cellular senescence is the development of dysfunctional telomeres^21^. These are generated not only in response to extensive cell proliferation but also as a consequence of genotoxic stresses that cause double-stranded DNA breaks in telomeric repeats^22,23^. Senescent cells are characterized by the expression of cyclin-dependent kinase inhibitors (CDKi), including p15INK4B, p16INK4A, and p21CDKN1A, as well as enhanced secretion of proinflammatory cytokines, and the expression of antiapoptotic genes, which makes them resistant to apoptosis^24^. Senescent cells additionally develop a greater abundance of lysosomal content, resulting in increased expression of lysosomal βGalactosidase^25^, also known as senescence-associated βGalactosidase (SA-βGal) in the context of cellular senescence. This hallmark of senescent cells allows their detection *in vitro* and *in vivo*, regardless of the senescence pathway activated^26^.

We previously reported a steep age-associated increase of SA-βGal expressing human peripheral blood CD8 T cells across all four T cell differentiation states and identified CD8 T_EM_ cells as a main component of the SA-ßGal expressing CD8 T cell pool detected in older individuals^8^. In this study, we further characterized the senescence state of SA-βGal-expressing CD8 T_EM_ cells, their functional capabilities, and their enrichment in aging, exposure to toxins, and disease. Our study offers novel and important insights into the functional impairment of senescent CD8 T cells, reveals their involvement in aging and disease, and opens new opportunities to predict outcomes of immunotherapies.

## Results

### Single-cell RNA sequencing identifies a transcriptionally unique SA-ßGal^high^ CD8 T_EM_ cell population that progressively expands with human aging

We previously demonstrated that CD8 T cells with high SA-ßGal activity exist across all four T cell differentiation states (T_N_, T_CM_, T_EM,_ T_EMRA_)^8,27^, classified by CCR7 and CD45RA expression^28^, suggesting that cellular senescence is decoupled from differentiation. To determine whether SA-ßGal identifies transcriptionally distinct populations within each of these differentiation states, we sorted approximately equal numbers of CD8 T cells with very low and very high SA-ßGal activity in each of the four T cell differentiation states and performed single-cell RNA sequencing (scRNAseq), retrieving 32,960 high-quality CD8 T cells for further analysis. Cells were isolated from 2 healthy human donors and sorted by fluorescence-activated cell sorting (FACS) using the fluorogenic ßGal substrate SPiDER-ßGal as previously described by us^8^ (**Figure S1A)**. Uniform Manifold Approximation and Projection (UMAP) embedding and clustering revealed eight transcriptionally distinct populations, three each within T_EMRA_ (EMRA1, EMRA2, EMRA3) and T_EM_ (EM1, EM2, EM3) populations, and one each within T_CM_ (CM) and T_N_ (N) populations (**Figure 1A-C**), as annotated based on a recent study^29^. While SA-ßGal^low^ and SA-ßGal^high^ cells were confined to a single cluster for the T_N_ and T_CM_ cell populations, SA-ßGal^high^ cells were significantly enriched in clusters EMRA2, EMRA3, and EM2 (**Figure 1B, D**). To reveal transcriptional differences between SA-ßGal^low^ and SA-ßGal^high^ cells that were masked due to T cell differentiation, we reclustered each of the four T cell differentiation states separately, which revealed that SA-ßGal^high^ CD8 T cells were transcriptionally distinct from SA-ßGal^low^ CD8 T cells in all four differentiation states, albeit to varying degrees.

**Figure 1.**
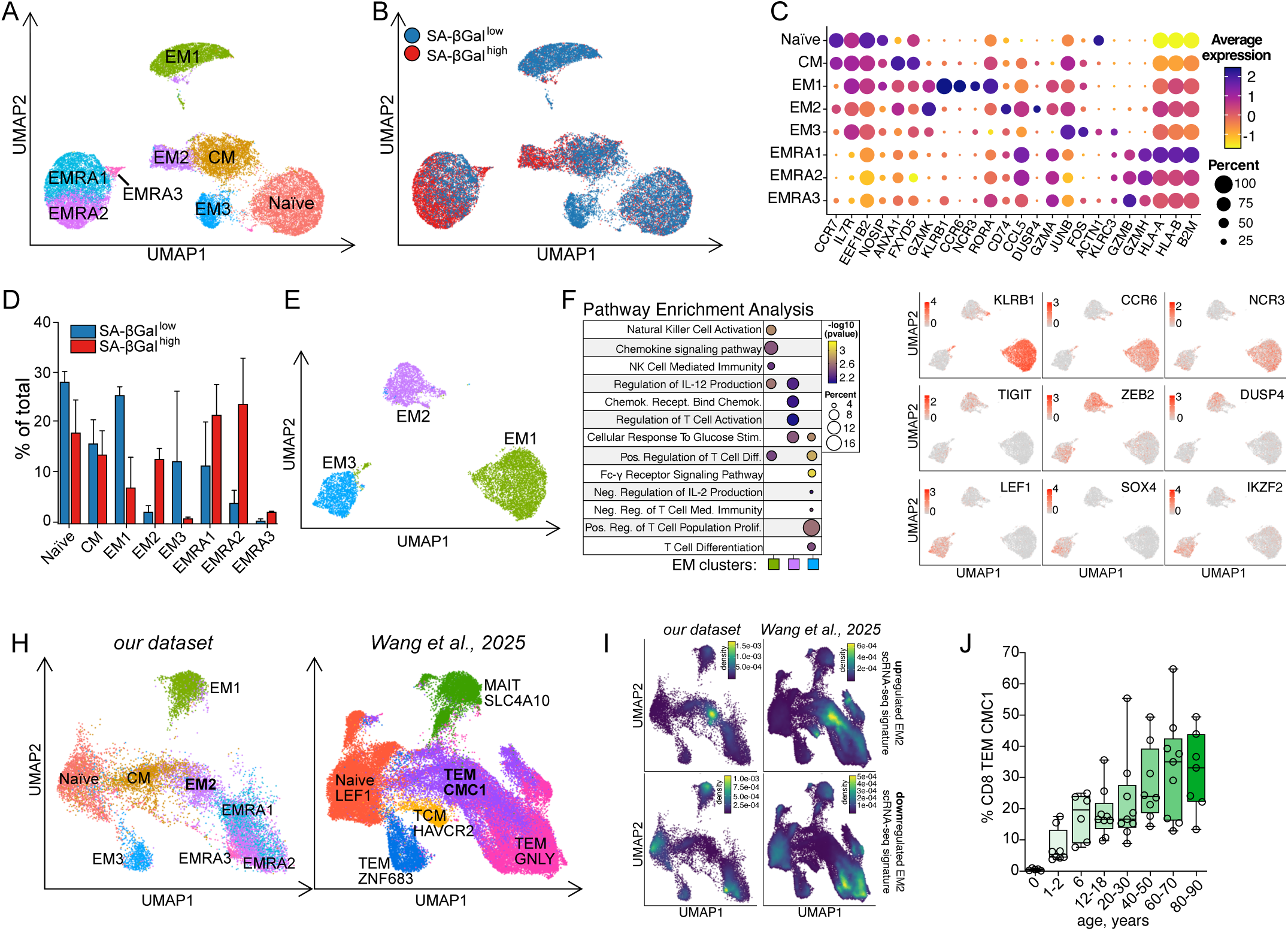
Single-cell RNA sequencing identifies a transcriptionally unique SA-ßGal^high^ CD8 T_EM_ cell population that progressively expands with human aging. **A.** UMAP projection of 32,960 FACS-sorted CD3+CD8 T cells from 2 healthy human donors, annotated by T-cell subset. **B**. Same UMAP as in (A), with cells colored based on SA-βGal sorting gate (blue: SA-βGal^low^; red: SA-βGal^high^). **C.** Dotplot of CD3+/CD8 T cell clusters identified in (A). Average expression levels and percentage of cells expressing the indicated marker genes are shown. **D**. Quantification of the percentage contribution of each cluster to the SA-βGal^low^ (blue) or SA-βGal^high^ (red) populations. Data are represented as mean +/- SD. **E**. UMAP projection of clusters EM1 (green), EM2 (purple), and EM3 (blue). F. Functional overrepresentation analysis map showing significant associations between indicated gene sets and EM1, EM2, and EM3 clusters. Circle fill is color-coded by false discovery rate (FDR)-corrected p-value from a hypergeometric distribution test. Circle size is proportional to the percent of genes in indicated gene sets. G. UMAP projections of T_EM_ clusters showing cluster-specific expression of indicated genes. **H.** UMAP projection of CD3+/CD8 T cells derived from our dataset and from *Wang et al., 2025*^38^. Cells are split by dataset as indicated. **I.** Gene set enrichment analyses using AUCell scoring, transformed into density scores, mapped onto UMAP projections for EM2-specific upregulated (top row) or downregulated (bottom row) genes from our dataset (left column), or that from *Wang et al., 2025*^38^ (right). **J.** Quantification of the percentages of cells in the TEM_CMC1 cluster out of the total CD8 T cell population in the indicated age groups, shown as a box- and whisker-plot (min to max). Each circle represents one donor.

Among the four CD8 T cell differentiation states, we observed the greatest transcriptional distinction within CD8 T_EM_ cell populations. UMAPs generated two distinct clusters enriched in SA-ßGal^low^ cells (EM1 and EM3) and one cluster that was almost entirely comprised of SA-ßGal^high^ cells (EM2) (**Figure 1D, E and S1B)**. Functional over-representation analysis on differentially expressed genes (DEGs) between the three T_EM_ clusters revealed enrichment for pathways such as “Natural Killer Cell Activation” and “Natural Killer Cell Mediated Immunity” in cluster EM1, “Regulation Of T Cell Activation” and “Chemokine Receptor Bind Chemokines” in cluster EM2, and “T Cell Differentiation” and “Positive Regulation Of Cell Population Proliferation” in cluster EM3 (**Figure 1F**). Cells within cluster EM1 expressed high levels of KLRB1, CCR6, and NCR3, thereby resembling innate-like cytolytic cells known as Mucosal Associated Invariant T cells (MAITs)^30–32^, while cells in cluster EM3 expressed LEF1, SOX4, and IKZF2 (Helios), which is indicative of stem-like, memory-oriented cells^33,34^. SA-ßGal^high^ cells in cluster EM2 expressed high levels of inhibitory molecules TIGIT and DUSP4, as well as the transcription factor ZEB2, which is consistent with a dysfunctional terminal effector program^35–37^ (**Figure 1G**).

To characterize potential age-associated changes to SA-ßGal^high^ cells in cluster EM2, we mined scRNAseq datasets from two studies that sequenced peripheral blood CD8 T cells collected from larger cohorts of healthy human donors spanning the entire human lifespan^29,38^. Following preprocessing and integration, we removed dataset-specific clusters and annotated cells based on our analysis above and that described in Wang et. Al 2025^38^ (**Figure S1C-D)**. Integration retained the transcriptional distinction between SA-ßGal^low^ and SA-ßGal^high^ cells, allowing us to identify these two populations within this new integrative model (**Figure S1E**). Our analysis demonstrated high concordance between the two datasets, as evident by the matching cell states among the corresponding clusters of each dataset, and revealed transcriptional similarities between the TEM CMC1 cluster and our EM2 cluster, enriched in SA-ßGal^high^ T_EM_ cells (**Figure 1H**). To determine whether SA-ßGal^high^ T_EM_ cells in cluster EM2 are identical to those in cluster TEM_CMC1, we generated EM2-specific gene signatures containing genes uniquely up- or downregulated in cluster EM2, compared to all other clusters in our scRNAseq dataset. Gene set enrichment analysis (GSEA) revealed that these signatures accurately discriminated only cells in clusters EM2 and TEM_CMC1 from all other cells, demonstrating that cell populations within the TEM_CMC1 and EM2 clusters are identical (**Figure 1I).** Significantly, consistent with our previous results that demonstrated an age-associated increase of SA-ßGal^high^ cells within the entire CD8 T cell population, this analysis revealed a substantial increase in the abundance of SA-ßGal^high^ CD8 T_EM_ cells within cluster TEM CMC1 with advancing age in healthy humans, reaching average levels of ∼35% of the total CD8 T cell population in advanced age (**Figure 1J)**.

Utilizing a similar approach, we also integrated our scRNAseq dataset with that from Lu et al.^29^, revealing the integrated cluster organization, annotation, and localization of the SA-ßGal low/high cell populations (**Figure S1 H-J**). Reclustering generated 16 clusters, each characterized by specific cell-type marker expression profiles (**Figure S1K)**, with cells in cluster 0 being identical to those in cluster EM2, as identified by GSEA using our EM2-specific SA-ßGal low/high gene signatures (**Figure S1M).** Similar to the TEM_CMC1 cell populations that increase in abundance with aging, we discovered that cluster 0 also progressively expands with advancing age in healthy humans (**Figure S1N)**. Overall, our data demonstrate that SA-ßGal activity identifies a transcriptionally unique SA-ßGal^high^ CD8 T_EM_ cell population that progressively expands with advancing age in healthy humans.

### SA-ßGal expressing CD8 T_EM_ cells are senescent

As CD8 T_EM_ cells are critically important for long-term and durable immunity, we sought to further characterize their senescence state, functional capabilities, and potential involvement in aging and disease. To collect SA-ßGal-low and high CD8 T_EM_ cells, we isolated CD8 T cells exclusively from healthy human donors who exhibited a clear bimodal distribution for SA-ßGal activity in the CD8 T_EM_ compartment, as determined by FACS (**Figure 2A and S2A**). From these donors, we consistently sorted the bottom fraction of cells in the βGal-low population (SA-ßGal^low^) and the top fraction of cells in the βGal-high population (SA-ßGal^high^) (**Figure 2A**), thereby ensuring the isolation of pure populations of SA-ßGal^low^ and SA-ßGal^high^ cells for all subsequent analyses.

**Figure 2.**
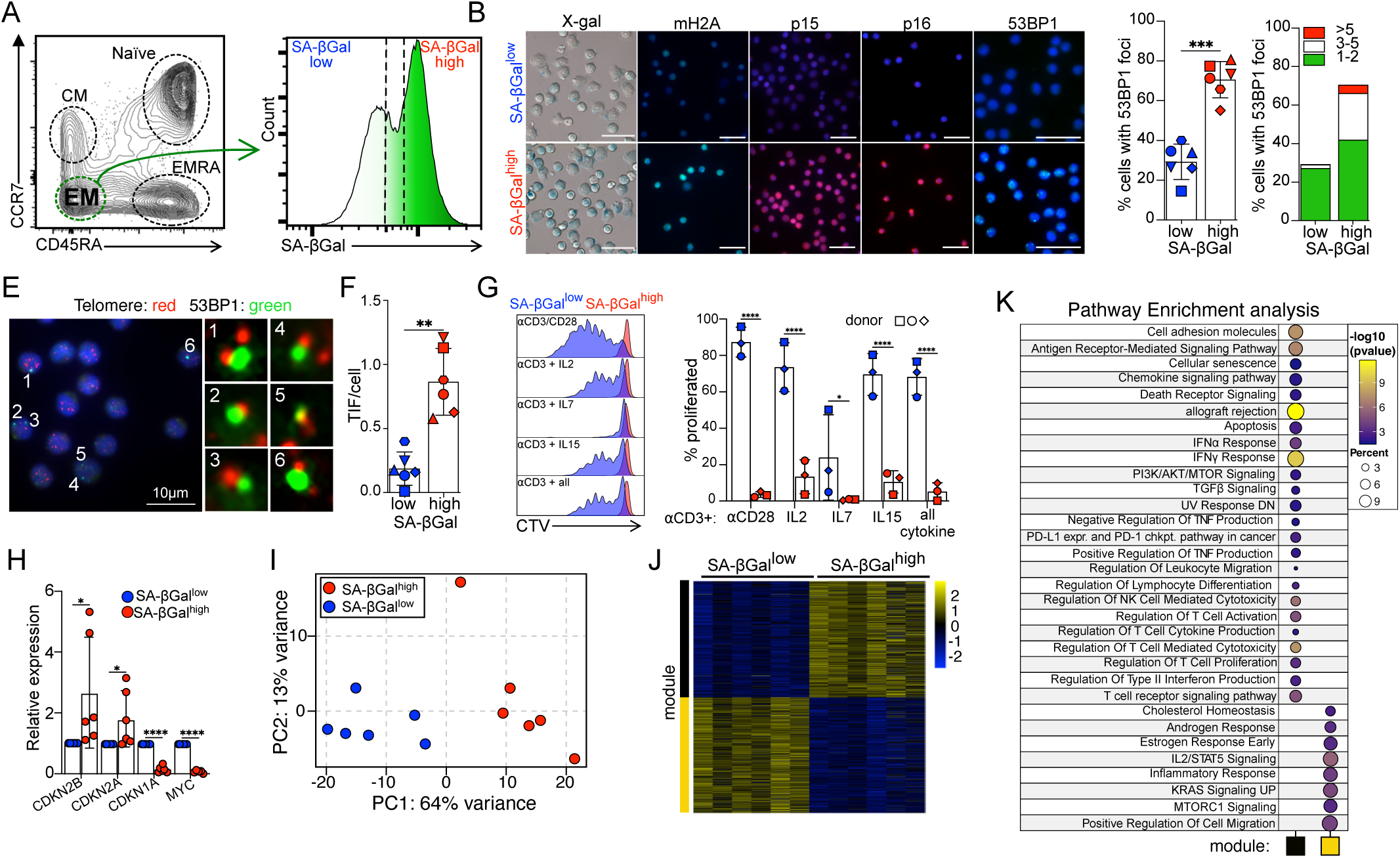
SA-ßGal^high^ CD8 T_EM_ cells are in a classical state of senescence. **A.** Representative FACS plots showing gating strategy to isolate CD8 T_EM_ cells based on absence of CCR7 and CD45RA expression (left) and to isolate SA-βGal^low^ and SA-βGal^high^ T_EM_ cells as indicated (right). **B**. Representative micrographs of SA-βGal^low^ (top row) and SA-βGal^high^ (bottom row) T_EM_ cells showing X-gal chromogenic staining or immunofluorescence analysis for mH2A, p15, p16, and 53BP1, as indicated. All immunostained cells were counterstained for DAPI (blue). White scale bars: 25μm. **C**. Quantification of the percentages of cells with 53BP1 foci in SA-βGal^low^ (blue) and SA-βGal^high^ (red) T_EM_ cells. Error bars: mean +/- SD. Shapes of data points correspond to unique donors. \*\*\**p* ≤0.001 by paired two-tailed t-test. **D**. Same quantification as in C with bar graphs stratified by the frequency of 53BP1 foci per cell (1-2: green, 3-5: white, >5: red). **E**. Sorted SA-βGal^high^ CD8 T_EM_ cells were simultaneously immunostained using antibodies against 53BP1 (green) and analyzed by FISH to detect telomeres (red). Blue: DAPI. Enlarged versions of the numbered DNA damage foci showing co-localization with telomeres are shown in the right micrographs. Scale bar: 10 μm. **F**. Quantification of mean TIF per cell in sorted CD8 T cells, as indicated. \*\**p* ≤0.01 by paired two-tailed t-test. **G**. Representative FACS profiles (left) and corresponding quantification (mean +/-SD) of Cell Trace Violet dilution assays in SA-βGal^low^ (blue) and SA-βGal^high^ (red) T_EM_ cells, stimulated for seven days with anti-CD3 antibodies plus indicated factors. Shapes of data points in the bar graphs correspond to unique donors. \**adj.p* ≤0.05,**** *adj.p* ≤0.0001 by one-way ANOVA and Holm-Šídák correction. **H**. RT-qPCR expression profiles for indicated transcripts in SA-βGal^low^ (blue) and SA-βGal^high^ (red) CD8 T_EM_ cells. n=6 for CDKN2B, CDKN2A, and CDKN1A; n=5 for MYC. Data are represented as mean +/- SD. \**p* ≤0.05,\*\*\*\**p* ≤0.0001 by paired one-tailed t-test. **I**. Principal component analysis (PCA) of transcriptomes (reads on exons) of SA-βGal^low^ (blue) and SA-βGal^high^ (red) CD8 T_EM_ cells. **J**. Weighted correlation network analysis (WGCNA) heatmap of color-coded modules showing 1128 differentially expressed genes (DEGs) between SA-βGal^low^ and SA-βGal^high^ CD8 T_EM_ cells; expression represented as a Z-score as indicated. **K**. Functional overrepresentation analysis map showing significant associations between indicated gene sets and modules. Circle fill is color-coded by false discovery rate (FDR)-corrected p-value from a hypergeometric distribution test, while circle size is proportional to the percentages of genes in indicated gene sets.

Unlike SA-ßGal^low^ CD8 T_EM_ cells that displayed discrete punctate foci of blue precipitates following staining with X-gal at pH=6, likely labeling their lysosomes and cytolytic granules, SA-ßGal^high^ CD8 T_EM_ cells exhibited robust X-gal staining throughout most of the cell body, demonstrating concordance between our fluorogenic and the more traditional chromogenic SA-ßGal detection assays^39^ (**Figures 2B)**. Consistent with a senescence state, SA-ßGal^high^ CD8T_EM_ cells expressed higher protein levels of p15, p16, and the senescence marker macroH2A^40^, compared to SA-ßGal^low^ cells, as detected by immunofluorescence analysis (**Figure 2B)**. Furthermore, approximately 70% of SA-ßGal^high^ CD8 T_EM_ cells displayed one or more discrete nuclear foci of 53BP1, a DNA damage response factor that localizes to the sites of double-stranded DNA breaks, while only ∼30% of SA-ßGa^low^ CD8 T_EM_ cells displayed such foci. (**Figure 2C-D)**. Many of these 53BP1 foci colocalized with telomeric repeats, demonstrating a high degree of telomere dysfunction-induced DNA damage foci (TIF) in SA-ßGal^high^ CD8 T_EM_ cells (**Figure 2E-F)**. To assess the proliferative capacity of CD8 T_EM_ cells with low and high SA-ßGal activity, we stimulated them either with anti-CD3/CD28 antibodies, to activate T cell receptor (TCR) signaling, or with various combinations of homeostatic cytokines IL2, Il7, and Il15 plus anti-CD3 antibodies, to account for potential proliferative defects that may arise from reduced expression of the co-stimulatory receptor CD28 by SA-ßGal^high^ CD8 T_EM_ cells (see **Figure S2B).** Proliferation was assessed by CellTrace Violet (CTV) dilution after 7 days of stimulation. Consistent with a classical senescence state, SA-ßGal^high^ CD8 T_EM_ cells were entirely impaired in their ability to proliferate, regardless of activating signals, while SA-ßGal^low^ CD8 T_EM_ cells proliferated extensively under most conditions tested (**Figure 2G).** To account for these proliferative defects, we measured expression of genes involved in cell cycle regulation and discovered significantly higher mRNA expression levels of the CDK inhibitors (CDKi) *CDKN2B* (p15) and *CDKN2A* (p16), as well as lower levels of *MYC*, in SA-ßGal^high^ CD8 T_EM_ cells compared to SA-ßGal^low^ CD8 T_EM_ cells, which is consistent with a senescence state (**Figure 2H**). Surprisingly, transcripts of the CDKi *CDKN1A* (p21) were largely undetectable in SA-ßGal^high^ CD8 T_EM_ cells, despite the presence of dysfunctional telomeres in these cells.

To characterize the transcriptomes of SA-ßGal^high^ CD8 T_EM_ cells in greater detail, we performed RNA-sequencing (RNAseq) in bulk of FACS-sorted SA-ßGal^low^ and SA-ßGal^high^ CD8 T_EM_ cells from 6 healthy human donors. Principal component analysis (PCA) revealed that transcriptomes of SA-ßGal^low^ and SA-ßGal^high^ CD8 T_EM_ cells were distinct, confirming our scRNAseq data (**Figure 2I**). Hierarchical clustering and self-organizing maps (SOM)^41^ corroborated these conclusions (**Figures S2C-D).** Weighted Gene Correlation Network Analysis (WGCNA) using the differentially expressed genes between SA-ßGal^low^ and SA-ßGal^high^ CD8 T_EM_ cells generated two modules, which partitioned genes upregulated in SA-ßGal^low^ (yellow module) and SA-ßGal^high^ (black module) (**Figure 2J**). Functional over-representation analysis revealed an enrichment for pathways including “cellular senescence”, “allograft rejection”, and inflammatory pathways such as “IFNα and IFNy response” in the black module, consistent with a senescence phenotype. In contrast, the yellow module, enriched for pathways including “Positive Regulation of Cell Migration”, “IL2/STAT5 Signaling”, and “MTORC1 signaling”, which is characteristic of functional CD8 T cells (**Figure 2K)**. Taken together, our results demonstrate that CD8 T_EM_ cells with high SA-ßGal activity are in a classical state of senescence characterized by increased X-gal staining, elevated expression levels of p15, p16, and macroH2A, telomere dysfunction, a stable proliferative arrest, and a transcriptome consistent with a senescence state.

### Senescent SA-ßGal^high^ CD8 T_EM_ cells develop a distinct yet reduced transcriptional output following stimulation

The lack of proliferation in response to various signals suggests that senescent CD8 T_EM_ cells are intrinsically impaired in their ability to respond to activating signals. To test this, we characterized the transcriptomes of unstimulated and anti-CD3/CD28-stimulated (16 hrs) non-senescent and senescent CD8 T_EM_ cells by RNA sequencing in bulk. Principal component analysis (PCA) and hierarchical clustering revealed significantly distinct transcriptomes between non-senescent and senescent cells, both with and without stimulation, demonstrating that senescent CD8 T_EM_ cells are transcriptionally distinct from non-senescent CD8 T_EM_ cells, both basally and upon activation (**Figure 3A** and **S3A)**. Self-organizing maps (SOM)^41^ corroborated these conclusions (**Figure 3B)**. Significantly, comparing DEGs between non-senescent and senescent CD8 T_EM_ cells revealed a ∼40% reduction in stimulation-induced DEGs in senescent CD8 T_EM_ cells compared to non-senescent CD8 T_EM_ cells (3,979 vs. 6,588 DEGs, respectively), demonstrating that the transcriptional output is dramatically reduced upon TCR-stimulation in senescent CD8 T_EM_ cells (**Figure 3C**). While 3,175 stimulation-induced DEGs were common to both non-senescent and senescent CD8 T_EM_ cells, they were largely modulated in a similar fashion, albeit at different magnitudes (compare low vs high in heatmaps of **Figures 3D** and **S3B**), further highlighting the altered transcriptional response to stimulation by senescent CD8 T_EM_ cells. Despite this overall reduced transcriptional output, TCR stimulation created 794 DEGs that were unique to the senescence state, overall demonstrating that senescent CD8 T_EM_ cells mount not only a blunted, but also a unique response to TCR stimulation.

**Figure 3:**
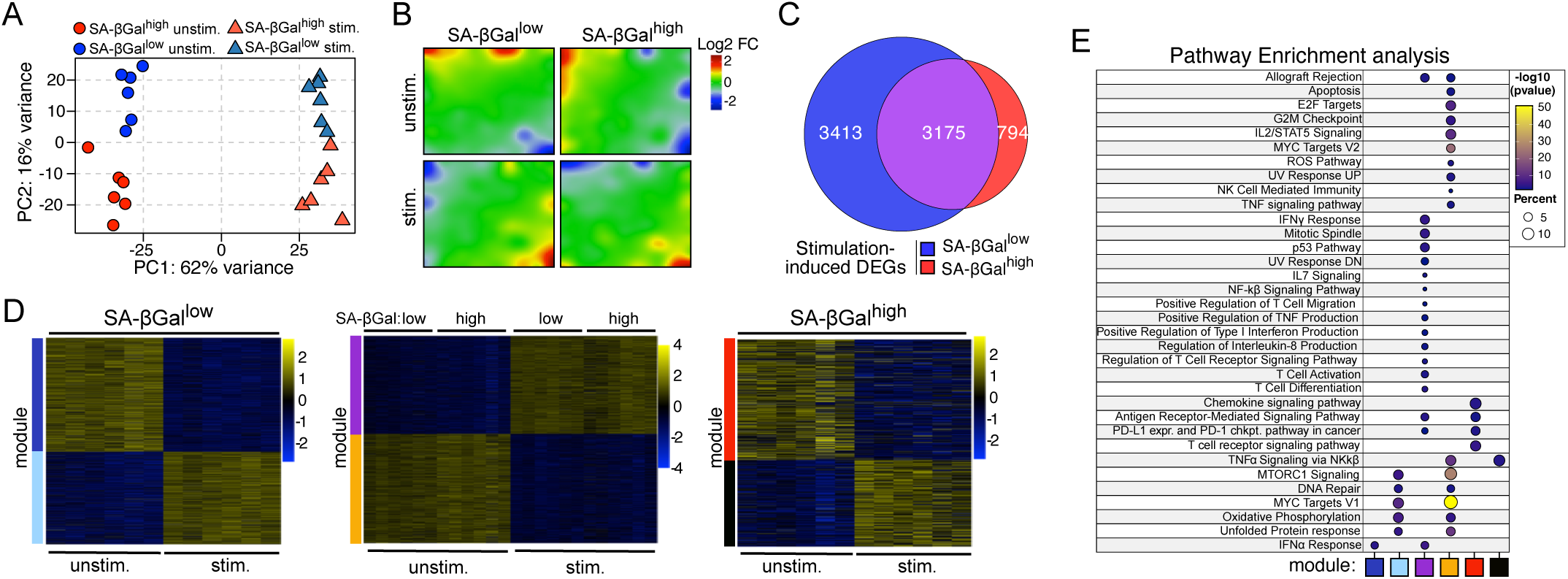
Senescent CD8 T_EM_ cells develop a distinct and blunted transcriptional output following stimulation. **A**. Principal component analysis (PCA) of transcriptomes (reads on exons) of SA-βGal^low^ (blue) and SA-βGal^high^ (red) T_EM_ cells. Circles: unstimulated, triangles: stimulated. **B**. Averaged self-organizing maps (SOMs) of transcriptomes from unstimulated and stimulated SA-βGal^low^ and SA-βGal^high^ T_EM_ cells as indicated. **C**. Venn diagram of stimulation-specific DEGs identified in SA-βGal^low^ (blue) and SA-βGal^high^ (red) CD8 T_EM_ cells; purple shading represents shared DEGs. **D**. WGCNA heatmaps of indicated color-coded modules from the SA-βGal^low^, shared, or SA-βGal^high^ CD8 T_EM_ stimulation-specific DEGs as indicated. **E**. Functional overrepresentation analysis map showing significant associations between indicated gene sets and modules. Circle fill is color-coded by false discovery rate (FDR)-corrected p-value from a hypergeometric distribution test, while circle size is proportional to the percentages of genes in indicated gene sets.

WGCNA clustering of stimulation-specific DEGs that were either shared or unique to non-senescent and senescent cells revealed 2 modules specific to non-senescent CD8 T_EM_ cells (blue, turquoise), 2 shared modules (purple and orange), and 2 modules specific to senescent CD8 T_EM_ cells (red and black). Whereas modules specific to genes overexpressed in non-senescent CD8 T_EM_ cells enriched for pathways that included “MYC targets”, those expressed in senescent CD8 T_EM_ cells enriched for “TNFα signaling via NFκB”. Stimulation-specific DEGs in the shared modules enriched for metabolic and cytokine secretion pathways. However, the magnitude of up- or downregulation for these pathways was noticeably different between non-senescent and senescent CD8 T_EM_ cells (**Figure S3C)**. In summary, our integrative transcriptomic analysis revealed that senescent CD8 T_EM_ cells mount a markedly reduced and unique response to TCR stimulation, characterized by the loss of proliferation-related programs and the gain of senescence-associated secretory pathways.

### Senescent SA-ßGal^high^ CD8 T_EM_ cells are functionally impaired

To test whether the senescence state impairs the functions of CD8 T_EM_ cells, we profiled their characteristics and stimulation-induced functional changes using a 19-color multiparameter spectral flow cytometry panel. Basally, the expression of markers that provide insights into the functional (TCF7, IL7R, CD28, PD1, CD16), cytolytic (Prf, GZMA, GZMB, GZMK, Lamp1), cytokine-producing (IL2, TNF, IFNγ), migratory, and signaling (CXCR1, CCR6, CCR9, CX3CR1, CXCR2, CXCR3) abilities were largely distinct between non-senescent and senescent CD8 T_EM_ cells (**Figures 4A and S4A**). These differences become even more apparent following 5h stimulation with either anti-CD3/CD28 antibodies or PMA/Ionomycin, which bypasses TCR signaling, causing several markers to change in opposite directions (TCF7, IL7R, Perforin, CXCR1, CCR6, CX3CR1) in non-senescent and senescent CD8 T_EM_ cells (**Figure S4A**). UMAP embeddings^42^ revealed that, overall, senescent CD8 T_EM_ cells remained largely unresponsive to TCR stimulation, although they developed a distinct response to stimulation with PMA/Ionomycin compared to non-senescent CD8 T_EM_ cells (**Figure 4B-C and S4A-C**). In addition, clusters of non-senescent CD8 T_EM_ cells overlapped only minimally with any of the senescent CD8 T_EM_ cell clusters, demonstrating that non-senescent CD8 T_EM_ cells do not develop a senescence phenotype following 5h stimulation (**Figure 4B-C and S4A-C**).

**Figure 4:**
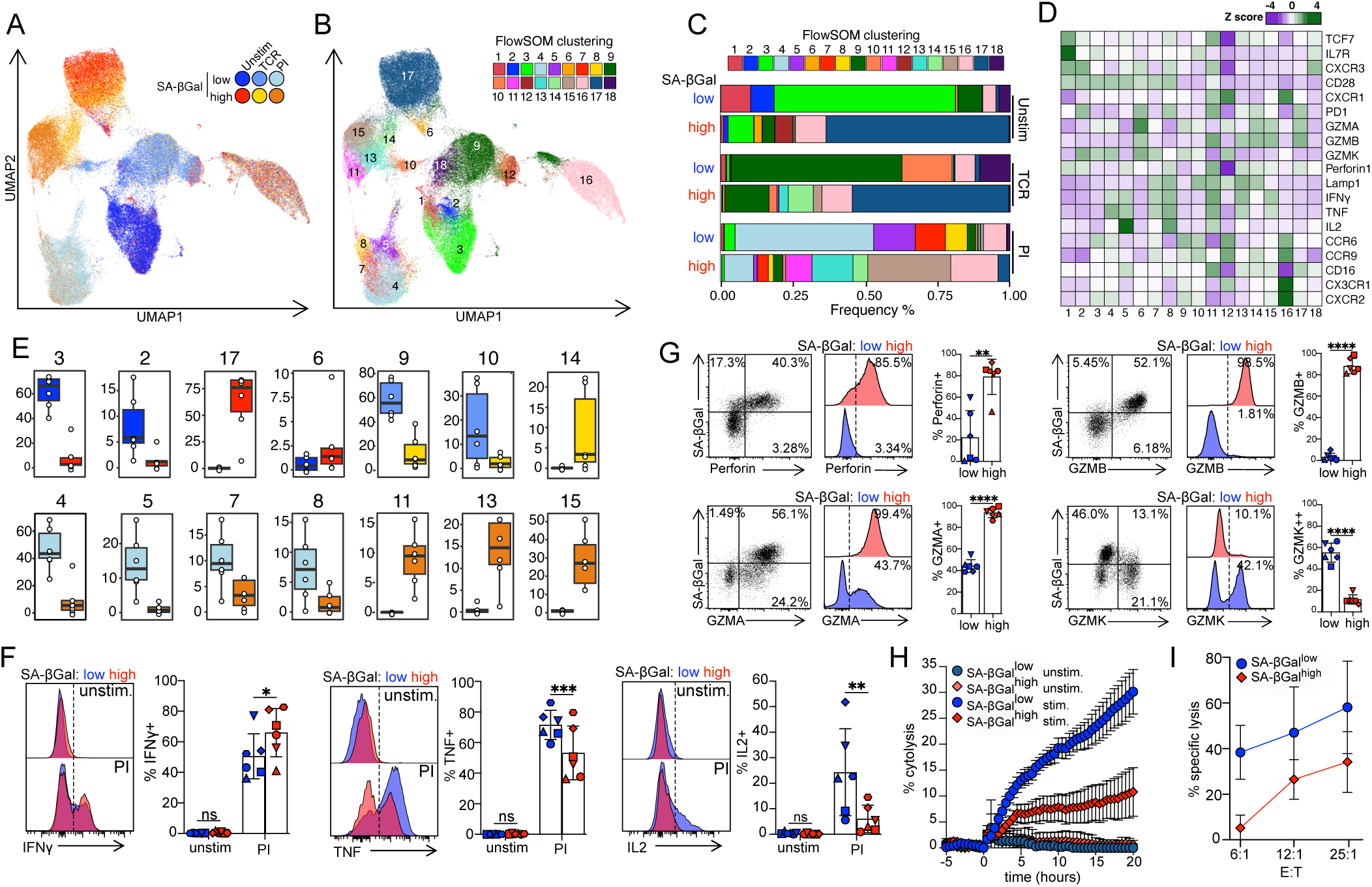
Senescent CD8 T_EM_ cells are functionally impaired. **A**. UMAP projection of FACS sorted SA-βGal^low^ (blue shading) and SA-βGal^high^ (red shading) CD8 T_EM_ cells that were either unstimulated (unstim.), or stimulated for 5 h with anti-CD3/CD28 beads to activate the T cell receptor (TCR) or with PMA/Ionomycin (PI) (n=6 donors). Embeddings are based on the expression pattern of 19 makers. **B**. Same UMAP as in (A), with cells colored by FlowSOM clusters. **C**. Sacked bar graphs showing the percentages of each FlowSOM cluster under indicated conditions. **D**. Expression pattern heatmaps (Z score) of indicated 19 markers across the 18 FlowSOM clusters. **E**. Box plots showing the % of sorted SA-βGal^low^ and SA-βGal^high^ CD8 T_EM_ cells within select FlowSOM clusters. Colors of boxes represent different stimulation conditions, as in A. Each circle represents a unique donor. **F**. Representative FACS dot plots, gating strategy, and quantification of Perforin, GzmB, GzmA, and GzmK from SA-βGal^low^ and SA-βGal^high^ CD8 T_EM_ cells of freshly isolated human CD8 T cells. Shapes of data points in the bar graphs correspond to unique donors. Error bars: +/- SD; ** *p* ≤0.01,\*\*\*\**p* ≤0.0001 by paired two-tailed t-test. **G**. Representative FACS histograms showing gating strategy and quantification (mean +/- SD) of indicated cytokines in unstimulated (unstim.) and 5-hour PI-stimulated FACS-sorted SA-βGal^low^ (blue) and SA-βGal^high^ (red) CD8 T_EM_ cells. Shapes of data points in the bar graphs correspond to unique donors. \**adj.p* ≤0.05, \*\**adj.p* ≤0.01,\*\*\**adj.p* ≤0.001 by two-way ANOVA and repeated measure Šídák correction. **H** Quantification of cytolysis of senescent human GM21 fibroblasts cocultured with unstimulated or 72h TCR pre-stimulated SA-βGal^low^ (blue) and SA-βGal^high^ (red) CD8 T_EM_ cells, as indicated, at an effector:target ratio of 25:1. T cells were added at hour 0. Error bars: SEM; n=5 for unstimulated and n=7 for stimulated effector cells. **I**. Percent cytolysis of Raji B cell lymphoma cells cocultured for 16 h with prestimulated SA-βGal^low^ (blue, circles) and SA-βGal^high^ (red, diamonds) CD8 T_EM_ cells at indicated effector:target ratios., \*\*\*\**p* ≤0.001 by two-way ANOVA with Holm-Šídák correction using a single pooled variance. Data points are shown as mean +/- SD, n=3.

FlowSOM^43^ identified 18 distinct clusters within these UMAPs, with non-senescent and unstimulated cells predominantly occupying cluster 3, characterized by high levels of stem-like markers, including TCF7, IL7R, and CD28, as well as the tryptase-like serine protease GZMK. In contrast, senescent CD8 T_EM_ cells were enriched in cluster 17, characterized by high expression levels of GZMA, GZMB, and PD1. Notably, a fraction of senescent CD8 T_EM_ subset expressed CD28 (see clusters 1, 2, 3, 6), while many non-senescent cells did not (see 9, 16, 17, 18), demonstrating that the absence of CD28 expression does not accurately characterize senescent SA-ßGal^high^ T_EM_ cells **(Figures 4C and S2B).** Significantly, while TCR stimulation caused an expansion of non-senescent T_EM_ cells in clusters 9, 10, and 18, characterized by LAMP1 externalization, IL-2 production, and upregulation of chemokine receptors CCR6 and CCR9, it caused only minor shifts in the cluster distribution of senescent T_EM_ cells (Figures **4C** **and S2B).** Similarly, stimulation with PMA/ionomycin resulted in the expansion of clusters 4, 5, 7, and 8 in non-senescent cells, and of clusters 11, 13, and 15 in senescent cells, revealing distinct stimulation-induced trajectories between these two cell states. PMA/Ionomycin stimulation caused both non-senescent and senescent CD8 T_EM_ cells to degranulate, externalize LAMP1, and produce IFN-γ and TNF. IL-2-producing cells, however, were predominantly confined to the non-senescent cell population, revealing a senescence-associated deficit in IL-2 production that goes beyond receptor signaling (**Figures 4D-E** and **Figures S4B-C**). To validate these results, we re-analyzed our FACS data using traditional gating and found that levels of IL-2 and TNF production were significantly reduced in senescent CD8 T_EM_ cells, while IFN-γ levels were increased, following PMA/Ionomycin stimulation (**Figure 4F**). Boolean gating on LAMP1-positive and negative cells expressing IFNy, TNF, and IL2+ cells revealed that non-senescent CD8 T_EM_ cells contained a higher fraction of polyfunctional T cells (IFNy+TNF+IL2+) and, on average, also greater numbers of degranulated polyfunctional LAMP1+ cells, compared to senescent cells, albeit at levels that did not reach statistical significance (**Figure S4D**). In contrast to non-senescent CD8 T_EM_ cells, senescent CD8 T_EM_ cells expressed low levels of GZMK and high levels of perforin, GZMA, and GZMB, suggesting that these cells are highly cytolytic (**Figure 4G**). Surprisingly, while protein expression levels of GZMB were high and those of GZMK were low in senescent SA-ßGal^high^ CD8 T_EM_ cells, their transcripts showed the opposite expression pattern in our scRNAseq data. This demonstrates that scRNAseq data do not reliably predict protein expression levels or the potential for T cell-mediated cytolysis.

Human CD8 T cells can target and eliminate senescent human fibroblasts in autologous and non-autologous cell targeting assays *ex vivo*^9,10^. To characterize the cytolytic activity of senescent CD8 T_EM_ cells, we co-cultured SA-ßGal^low^ and SA-ßGal^high^ CD8 T_EM_ cells, either unstimulated or pre-stimulated with anti-CD3/CD28 plus IL2, with senescent human fibroblasts for 20 h in xCELLigence RTCA microplates and monitored cell lysis in real time. While unstimulated CD8 T_EM_ cells had negligible cytolytic activity, regardless of whether they were senescent or not, pre-stimulation of the TCR caused non-senescent CD8 T_EM_ cells to effectively target and lyse senescent human fibroblasts under these conditions (**Figure 4H**). Surprisingly, senescent CD8 T_EM_ cells were significantly impaired in their cytolytic activity towards senescent fibroblasts, despite degranulating (**Figure S4E**) and expressing high levels of cytolytic molecules, prior to and three days after stimulation (**Figures 4G-H and 4F**). Consistent with these results, we discovered that the cytolytic activity of senescent CD8 T_EM_ cells towards human B-cell lymphoma cells (Raji) was similarly impaired compared to that of non-senescent CD8 T_EM_ cells at all effector-to-target ratios analyzed (**Figure 4H).** Collectively, our findings reveal that senescent CD8 T_EM_ cells lack polyfunctionality and harbor cell-intrinsic defects in cytotoxicity, thereby failing to lyse senescent or cancer cells despite harboring abundant cytolytic granule proteins.

### The senescent CD8 T_EM_ cell gene signature is enriched in aging, exposure to toxins, and age-related diseases

Defective mouse T cells, as well as senescent and aged mouse splenocytes, promote the accumulation of senescent non-lymphoid cells, age-related disorders, and aging in mice^5–7^. As senescent CD8 T_EM_ cells are dysfunctional and increase in abundance with aging in humans, as revealed in this and our previous studies^8^, we investigated their involvement in aging, age-associated disorders, exposure to toxins, and cancer. To this end, we generated gene signatures for senescent and non-senescent CD8 T_EM_ cells from freshly isolated, unstimulated CD8 T_EM_ cells with high and low SA-ßGal activity, respectively, and mined published transcriptomic datasets of human peripheral blood CD8 T cells isolated from donors in various study cohorts. Predictably, our senescent gene signature was enriched in circulating CD8 T_EM_ cells of healthy older adults (ages 62-75) compared to younger adults (ages 23-28)^44^, in agreement with our scRNAseq data (see Figure 1J and S1N) and our previous studies^8^. Significantly, our senescent gene signature was also enriched in CD8 T cells from smokers, compared to non-smokers^45^, as well as in CD8 T cells from patients with rheumatoid arthritis^46^ and advanced-stage melanoma^47^, compared to healthy individuals (**Figure 5A-D)**. Conversely, our non-senescent CD8 T_EM_ cell gene signature was enriched in transcriptomes from CD8 T cells isolated from younger and healthy individuals from the same cohorts (**Figure S5A-D)**, overall highlighting the utility of these gene signatures in facilitating our understanding of the involvement of senescent CD8 T_EM_ cells in aging and disease.

**Figure 5.**
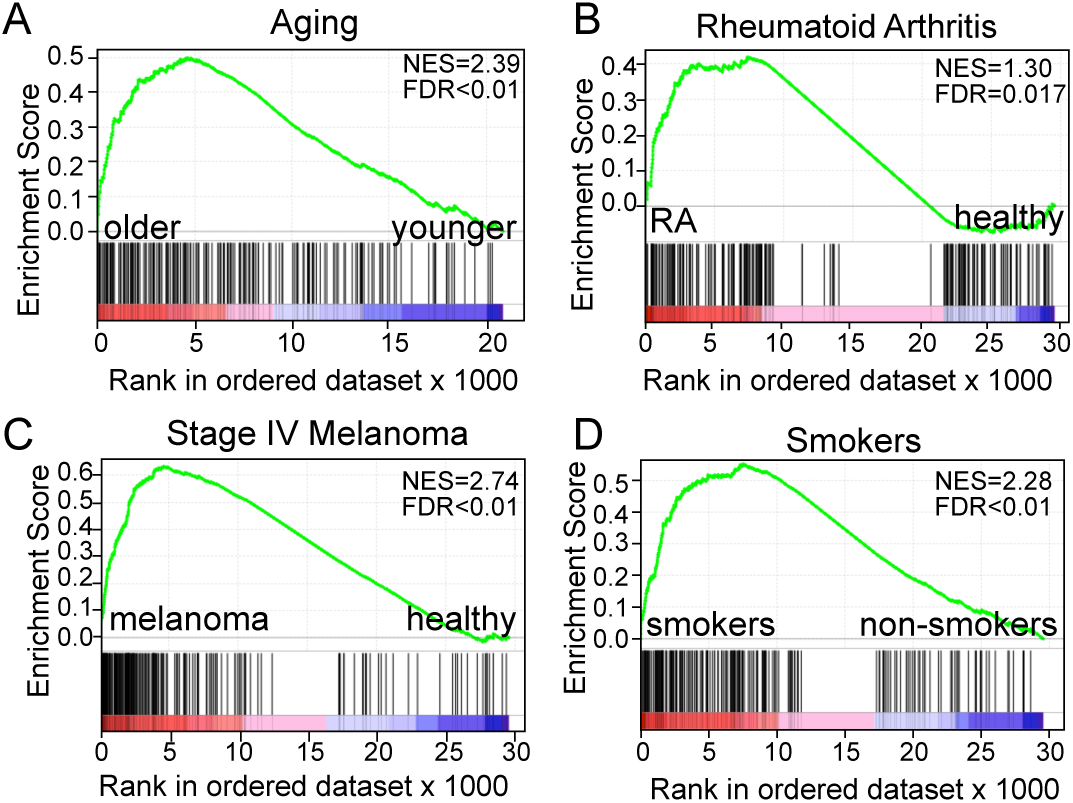
A senescent CD8 T_EM_ cell gene signature is enriched in CD8 T cells during aging, exposure to toxins, and age-related diseases. Gene set enrichment analyses (GSEA) showing normalized enrichment score (NES) plots and false discovery rate (FDR) values for our senescent CD8 T_EM_ cell gene signature that enriches in normalized and ranked transcriptiomes of peripheral blood CD8 T_EM_ cells isolated from older healthy human donors (**A**), patients with rheumatoid arthritis (RA) (**B**), patients with stage IV melanoma, and from peripheral blood CD8 T cells of smokers (**D**).

### CD8 T_EM_ cell senescence is distinct from exhaustion and determines responsiveness to anti-PD-1 immunotherapy

Similar to cellular senescence, exhaustion is a dysfunctional T cell state characterized by diminished proliferative capacity and effector function^19^. To investigate the relationship between T cell senescence and exhaustion, we first measured PD-1 expression across the four CD8 T cell differentiation states, revealing the greatest abundance of PD-1-expressing cells in T_EM_ CD8 T cell populations (**Figure 6A**). Surprisingly, we could not detect a clear correlation between PD-1 expression and SA-βGal activity in CD8 T_EM_ cells, as all four populations (SA-βGal^high^/PD-1^+^, SA-βGal^low^/PD-1^+^, SA-βGal^high^/PD-1^-^, and SA-βGal^low^/PD-1^-^) coexisted in our study cohort of 11 healthy human donors (**Figure 6B-C**). Although PD-1-expressing cells were, on average, more frequently detected in CD8 T_EM_ populations that were in a state of senescence (see **Figure S6A**), the relative distribution of the four PD-1/SA-ßGal populations varied dramatically between healthy donors. While some donors expressed PD-1 almost exclusively on either SA-ßGal^low^ or SA-ßGal^high^ CD8 T_EM_ cells (for example, see donors 3 and 9 in **Figure 6C**), most other donors displayed a more balanced distribution of these four populations (**Figure 6C**). This suggests that T cell exhaustion and senescence represent two distinct cell states.

**Figure 6.**
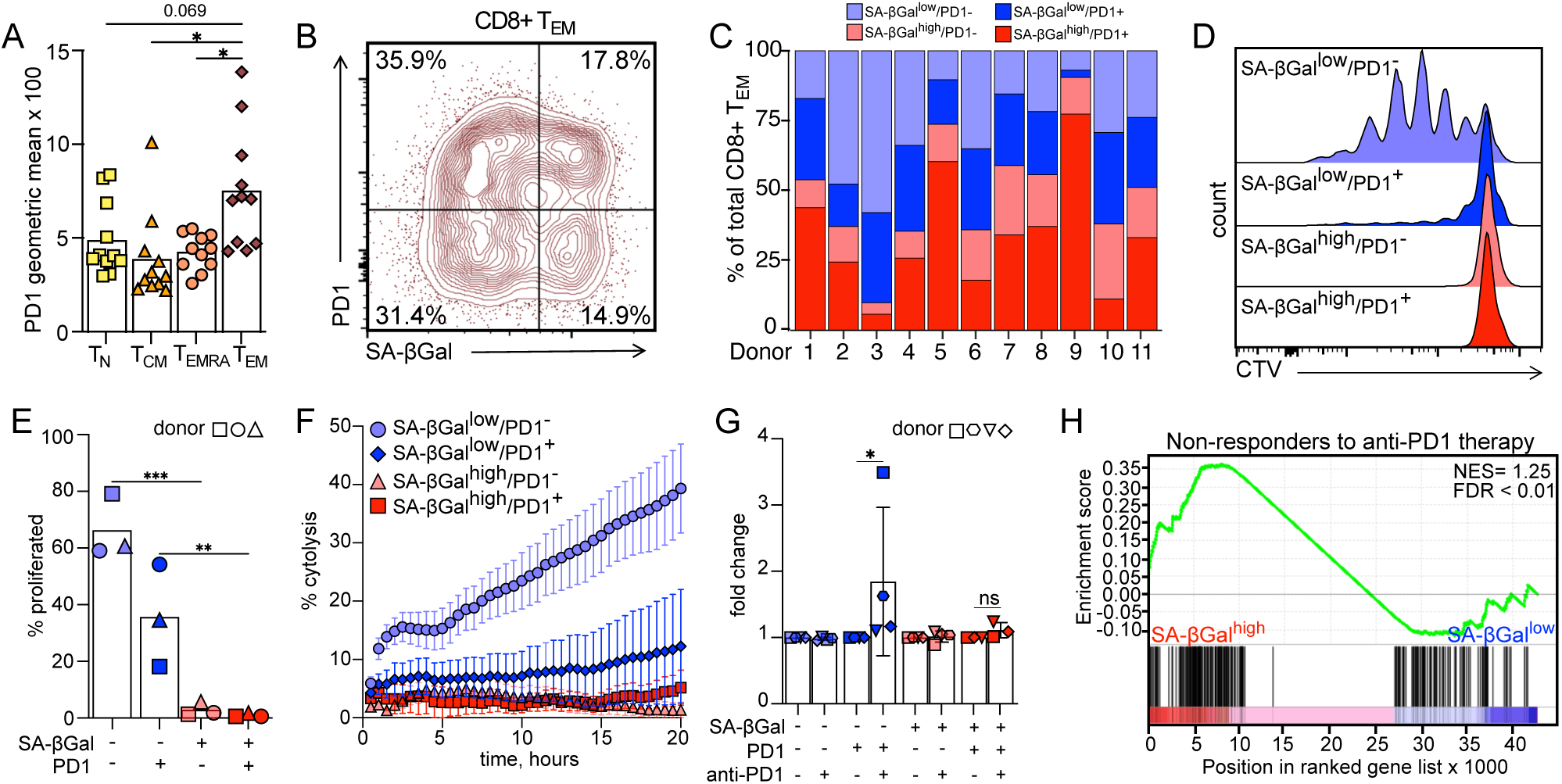
CD8 T_EM_ cell senescence is distinct from exhaustion and determines responsiveness to anti-PD-1 immunotherapy. **A**. Quantification of PD1 expression as geometric mean intensity across indicated CD8 T cell differentiation states. Bar graphs represent mean values; n=11. \**adj.p* ≤0.05 by one-way ANOVA and repeated measure Bonferroni correction. **B**. Representative FACS contour plot and gating strategy to quantify PD1- and SA-βGal expressing populations within the CD8 T_EM_ subset. **C**. Stacked bar graph showing the proportion of indicated populations in 11 healthy human donors. **D**. Representative FACS profiles of Cell Trace Violet dilution assays in FACS sorted CD8 T_EM_ populations, as indicated, that were TCR stimulated for five days. **E**. Quantification of indicated cell populations, color coded as in D, that had undergone at least one cell division; bar graphs represent mean; shapes of data points in the bar graphs correspond to unique donors. \**adj.p* ≤0.05, \*\**adj.p* ≤0.01 by one-way ANOVA and repeated measure Bonferroni correction using a single pooled variance. **F**. Quantification of cytolysis of senescent human GM21 fibroblasts cocultured with unstimulated or 72h TCR pre-stimulated CD8 T_EM_ cell populations, as indicated, at an effector:target ratio of 25:1. T cells were added at hour 0. Error bars: SEM; n=4 for SA-βGal^high^ PD1+; n=5 for all other populations. **G**. Fold change of relative cytolysis of indicated CD8 T_EM_ populations following anti-PD1 (Nivolumab) treatment. Shapes of data points in the bar graphs correspond to unique donors. Bar graphs are represented as mean +/- SD. \**p* ≤0.05 by paired two-tailed t-test. **H**. GSEA showing the NES plots and FDR values for a peripheral blood CD8 T cell gene signature from non-responders to anti-PD1 therapy that enriches in normalized and ranked transcriptomes from senescent SA-βGal^high^ CD8 T_EM_ cells.

Proliferation following TCR stimulation was greatest in SA-ßGal^low^/PD-1^-^ populations, followed by SA-ßGal^low^/PD-1^+^ cells, revealing proliferative defects of PD-1-expressing CD8 T cells, consistent with previous findings^48,49^. In contrast, senescent SA-ßGal-expressing CD8 T_EM_ cells failed to proliferate irrespective of PD-1 expression, confirming that the senescence state causes a stable proliferative arrest of CD8 T_EM_ cells (**Figure 6D-E)**. Similarly, we discovered the greatest cytolytic activity of CD8 T_EM_ cells towards senescent human PD-L1-expressing fibroblasts (**Figure S6B**) in SA-ßGal^low^/PD-1^-^ populations, followed by SA-ßGal^low^/PD-1^+^ cells, while senescent SA-βGal-expressing CD8 T_EM_ cells essentially failed to target and lyse senescent fibroblasts, regardless of whether they expressed PD-1 or not (**Figure 6F**). Overall, our data therefore demonstrate that the senescence state of CD8 T_EM_ cells is distinct from and dominant over CD8 T cell exhaustion.

Although PD-1-expressing CD8 T cells are impaired in their ability to lyse target cells expressing PD-L1, blocking this receptor-ligand interaction through ICB using anti-PD-1 antibodies can reverse their functional impairment and activate their cytolytic activity^19^. To test which of the four SA-ßGal/PD-1 populations respond to ICB, we repeated the cytolysis experiments with and without the therapeutic anti-PD-1 antibody Nivolumab, using PD-L1-expressing senescent human fibroblasts as target cells (see **Figure S6A)**. Significantly, while cytolytic activity of SA-ßGal^low^/PD-1^-^, SA-ßGal^high^/PD-1^-^, and SA-ßGal^high^/PD-1^+^ cells remained unchanged in the presence of Nivolumab, SA-ßGal^low^/PD-1^+^ cells responded to ICB and developed a ∼2 fold increase in cytolytic activity towards senescent cells in the presence of anti-PD-1 antibodies (**Figure 6G)**. Our data therefore demonstrate that the senescence state of CD8 T_EM_ cells restricts their ability to respond to ICB. Consistent with this conclusion, we found that the gene signatures of CD8 T cells from non-responders to ICB therapy in melanoma^50^ were highly enriched in senescent SA-ßGal^high^, but not non-senescent SA-ßGal^low^ CD8 T_EM_ cells (**Figure 6H)**, revealing an enrichment of senescent CD8 T_EM_ cells in melanoma patients refractory to ICB therapy.

## Discussion

As human CD8 T cells differentiate into the T_EMRA_ end-state, the majority of them lose expression of co-stimulatory receptors CD27 and CD28, exhibit KLRG1 and CD57 on their cell surface, upregulate the senescence markers p21 and p16, and show reduced proliferative capacity^16^. Despite displaying these senescence features, however, many terminally differentiated CD27^-^/CD28^-^ T_EMRA_ cells retain high cytolytic activity, which is mediated by NK-cell receptors and independent of TCR signaling^15^. Based on this, it has been proposed that senescent CD8 T cells develop innate immune cell-like cytolytic function, enabling them to continue targeting damaged, infected, and cancer cells in a TCR-independent manner^16^. In this study, we demonstrate that a substantial proportion of CD8 T cells with classical features of senescence also develop outside of the T_EMRA_ differentiation state. This senescent CD8 T cell population develops in the effector memory (EM) compartment and is characterized by high SA-ßGal activity, elevated levels of p15^INK4b^, p16^INK4a^, and macroH2A, and is independent of CD28 expression. In contrast to CD27^-^/CD28^-^ T_EMRA_ cells, senescent SA-ßGal^high^ CD8 T_EM_ cells completely lack the ability to proliferate, regardless of stimulating signals, and are significantly impaired in their cytolytic activity towards senescent and cancer cells, despite retaining high levels of cytolytic molecules. Our data therefore reveal a novel, previously unrecognized population of senescent and dysfunctional CD8 T cells, detectable based on high SA-ßGal activity.

The functional activity of CD8 T cells is largely defined by their ability to release cytokines that induce proliferation, modulate inflammation, and mediate cytolysis, as well as by their ability to release cytolytic molecules through a process termed degranulation, which enables them to lyse target cells. Our data reveal that senescent SA-ßGal^high^ CD8 T_EM_ cells are significantly impaired in most of these functions, including their ability to proliferate, produce cytokines IL-2 and TNF, as well as their cytolytic activity towards senescent cells and cancer cells following stimulation. While most senescent SA-ßGal^high^ CD8 T_EM_ cells lacked expression of co-stimulatory receptor CD28, which could partially account for their unresponsiveness to stimulation with anti-CD3/CD28 antibodies, senescent SA-ßGal^high^ CD8 T_EM_ cells also exhibited significant defects following stimulation with cytokines as well as with PMA/Ionomycin, which activate CD8 T cells independent of T cell receptor-signaling. This demonstrates that senescent SA-ßGal^high^ CD8 T_EM_ cells are functionally impaired irrespective of CD28 expression. The lack of cytolytic activity of senescent SA-ßGal^high^ CD8 T_EM_ cells, however, is surprising, given that they express high levels of perforin and granzymes basally, while also retaining the ability to degranulate. Although we currently do not know the causes for their cytolytic defects, it is possible that senescent SA-ßGal^high^ CD8 T_EM_ cells fail to polarize their lytic granules towards the immunological synapse, as has been reported for aged and defective NK cells^51^. The absence of cytolytic activity by senescent SA-ßGal^high^ CD8 T_EM_ cells is also in contrast to senescent-like CD27^-^/CD28^-^ T_EMRA_ cells, reported to express various NK-cell receptors and exhibit high cytolytic activity towards cells expressing NKG2D ligands^15^. It is therefore likely that senescent and dysfunctional SA-ßGal^high^ CD8 T_EM_ cells resemble terminally exhausted human effector CD8 T cells described in chronic viral infections and characterized by limited polyfunctionality, proliferative capacity, and cytolytic activity^36,52^.

Single-cell RNA sequencing revealed three transcriptionally distinct CD8 T_EM_ cell populations (EM1-3), previously annotated based on known marker genes and validated using high-dimensional flow cytometry^29^. While CD8 T cells in cluster EM1 expressed transcripts consistent with those of mucosal-associated invariant T (MAIT) cells with tissue homing properties^53^, those in cluster EM3 upregulated genes characteristic of committed effector memory cells primed for activation^34^. In contrast, cells in cluster EM2, which was almost exclusively composed of senescent SA-ßGal^high^ CD8 T_EM_ cells, expressed genes for inhibitory molecules^37,54^, including TIGIT and DUSP4, as well as genes that characterize previously activated CD8 T cells^54,55^, including the HLR-DR class II molecules, and the invariant chain CD74, suggesting that CD8 T_EM_ cells entered a senescence state after they had been activated, proliferated, and expanded. Consistent with this interpretation, our analysis revealed a high degree of DNA damage and telomere dysfunction in senescent CD8 T_EM_ cells, features of cells that have entered a state of replicative senescence following a period of proliferation. Cells in cluster EM2 also expressed GZMK, but not GZMB transcripts, which defines clonal, pro-inflammatory CD8 T cell populations enriched in older humans^56^. By integrating our data with that from studies that sequenced CD8 T cells from larger cohorts of healthy aging humans, we further discovered that senescent SA-ßGal^high^ CD8 T_EM_ cells are transcriptionally analogous to CD8_TEM_CMC1 cells, a highly clonal CD8 T cell population that progressively increases in abundance with aging in healthy humans, reaching average levels of ∼35% of the total peripheral CD8 T cell population in advanced age^38^. Overall, our data therefore reveal that senescent SA-ßGal^high^ CD8 T_EM_ cells comprise a clonally expanded and dysfunctional cell population that displays features of replicative senescence and progressively increases in abundance with advancing age in healthy humans.

Since human CD8 T cells can target and eliminate senescent human non-lymphoid cells *ex vivo* and *in vivo,* as shown previously^9–13^ and in this study, it is likely that they are a component of the immune surveillance system responsible for clearing senescent cells that accumulate with aging systemically. Supporting this are studies demonstrating that perforin knockout mice with defective T and NK cells develop more senescent cells systemically, exhibit higher rates of age-related disorders, and have shorter lifespans compared to control animals^5^. The absence of cytolytic activity towards senescent fibroblasts and cancer cells by senescent SA-ßGal^high^ CD8 T_EM_ cells *ex vivo*, as demonstrated in this study, suggests that they are also unable to target and eliminate these cells *in vivo*. The age-associated increase in senescent SA-ßGal^high^ CD8 T_EM_ cells in healthy human donors may therefore increasingly impede efficient clearance of senescent cells with age systemically and lead to the development of age-associated disorders and diseases. Consistent with this interpretation, our data also revealed a significant enrichment of CD8 T cells with gene signatures of senescent SA-ßGal^high^ CD8 T_EM_ cells in patients with rheumatoid arthritis and melanoma. As our study did not establish causality, however, we currently do not know whether the enrichment of senescent SA-ßGal^high^ CD8 T_EM_ cells contributes to the development of these conditions, or whether the inflammatory environment associated with these disorders accelerates the development of senescent SA-ßGal^high^ CD8 T_EM_ cells in the periphery. Future studies aimed at establishing causality will provide further insights into the potentially damaging effects of senescent CD8 T cells on the organism.

Immune cell senescence and exhaustion are two dysfunctional cell states that have been shown to impede the efficient clearance of cancer cells^19^, as well as senescent cells that accumulate systemically during aging^12,57^. Although they are generally considered distinct cell states, the degree to which these dysfunctional states coexist and affect each other remains unclear. We previously demonstrated a lack of correlation between SA-ßGal levels and PD-1 expression in human peripheral blood CD8 T cells, suggesting that cellular senescence is decoupled from T-cell exhaustion^8^. In this study, we also failed to detect a correlation between exhaustion and senescence in CD8 T_EM_ cells, despite this population developing the largest fraction of PD-1-expressing cells within the four CD8 T cell differentiation states^58^. CD8 T_EM_ cell senescence and exhaustion, however, are not mutually exclusive cell states and can coexist within the same cell. We discovered that various fractions of both non-senescent and senescent CD8 T_EM_ cells express PD1, albeit at ratios that depend on the individual blood donor. Significantly, only non-senescent, PD-1-expressing cells responded to ICB, whereas senescent, PD-1-expressing cells did not, demonstrating that the senescence state limits the ability of PD-1-expressing CD8 cells to respond to ICB. This is analogous to the phenotype exhibited by terminally exhausted (Texh-term) CD8 T cells, a T cell state characterized by low TCF1 expression levels, low proliferative activity, and unresponsiveness to ICB^20^. While SA-ßGal^high^ CD8 T_EM_ cells share several features with Texh-term cells, including low expression levels of TCF1, whether Texh-term cells are in a state of senescence remains to be established. However, given that CD8 T cell gene signatures from non-responders to ICB are enriched in transcriptomes from senescent SA-ßGal^high^ CD8 T_EM_ cells, it is possible that senescent SA-ßGal^high^ CD8 T_EM_ cells contribute to the failure of cancer patients to respond to ICB therapy.

In this study, we identified a novel, and dysfunctional population of senescent SA-ßGal^high^ CD8 T_EM_ cells that exhibit classical features of cellular senescence and increase in abundance with advancing age in healthy humans. Their functional impairment and enrichment in aging and disease support the model that this senescent CD8 T_EM_ cell population contributes to age-associated systemic accumulation of senescent cells, potentially due to its failure to target and eliminate senescent cells in tissues. An enrichment of our senescent SA-ßGal^high^ T_EM_ gene signature in aging and disease, however, also suggests that gene signatures of senescent CD8 T cells may prove useful as biomarkers of immunological health, thereby allowing risk assessment for developing age-associated disorders and diseases. As our study also revealed the involvement of senescent SA-ßGal^high^ CD8 T_EM_ cells in the failure of patients to respond to ICB, it suggests that assessing the CD8 T cell senescence status of such patients might prove to be an important clinical tool for guiding ICB-based treatment decisions.

## Resource Availability

### Lead contact

For additional details or access to relevant materials, please reach out to the lead contact: Utz Herbig

### Materials Availability

This study did not generate new unique reagents.

### Data and Code Availability

Raw and processed data and the code used to generate the results and figures in this study will be deposited in the Gene Expression Omnibus, Sequence Read Archive and Github.

## ACKNOWLEDGMENTS

This research was supported by grants from the National Institute on Aging of the National Institutes of Health (R21AG091016) to UH, (R21AG067368) to UH and PFB, and a grant from the National Institute of General Medical Sciences of the National Institutes of Health (R35GM155447) to R.I.M-Z. The content is solely the responsibility of the authors and does not necessarily represent the official views of the National Institutes of Health.

## Author contributions

U.H and P.S.T conceived the study and designed experiments. P.F-B facilitated donor recruitment and blood collections. E.A, N.A, L.G-M, drew blood from human donors. P.S.T, E.A, N.A isolated PBMCs. S.S. provided insights and guidance in multiparameter flow cytometry, and FACS isolated SA-βGal low and high T_EM_ cells with P.S.T. G.Y provided insights on T cell biology. P.S.T conducted all experiments, with help from E.A in imaging and quantifying TIF per cell and N.A in acquiring CTV proliferation. P.S.T analyzed all RNA-sequencing data with guidance from R.I.M-Z. P.S.T and U.H assembled the figures and wrote the manuscript with input from all authors.

## Declaration of interest

The authors declare no competing interests.

## Experimental model and study participants details

The Institutional Review Board of New Jersey Medical School approved this study (ID Pro2020000006, approved May 14, 2024). Peripheral blood mononuclear cells (PBMCs) were obtained from the blood of consenting healthy donors collected in heparinized tubes (BD Biosciences, Franklin Lakes, NJ) or Leukopacks were commercially obtained from the New York Blood Center.

## Method details

### Peripheral blood mononuclear cell (PBMC) isolation and CD8 T cell enrichment

PBMCs were isolated using density gradient centrifugation with Lymphocyte Separation Medium (Corning, Manassas, VA). Briefly, heparinized blood was diluted 1:1 with Hank’s Balanced Salt Solution (HBSS) (Corning, Manassas, VA), layered over lymphocyte separation medium at a 2:1 ratio, and centrifuged at 400g for 30 minutes with reduced deceleration (deceleration factor of 2). The buffy coat, containing PBMCs, was carefully harvested and further processed. CD8 T cells were enriched from PBMCs using negative selection with the EasySep™ Human CD8 T Cell Isolation Kit (StemCell Technologies, Cambridge, MA), following the manufacturer’s protocol. The purity of the isolated CD8 T cells was assessed via flow cytometry by staining for surface markers PerCP/Cyanine7-CD3 (HIT3a), and PE-CD8 (RPA-T8), (BioLegend, San Diego, CA). Briefly, all surface staining was conducted at 4°C for 20 minutes in 1x PBS + 2% FCS, while protected from light. Samples were subsequently washed 1xPBS+ 2% FCS and kept on ice before being acquired on the BD LSRFortessa X-20 (BD Biosciences, Franklin Lakes, NJ). Samples consistently achieved >85% CD3+CD8 double-positive cells.

### Surface staining, SA-βGal staining, and FACS

Isolated and enriched CD8 T cells were resuspended in RPMI-1640 medium containing 10% fetal bovine serum (FBS, Atlanta Biologicals) and 1× penicillin/streptomycin (Corning, Manassas, VA). The Cellular Senescence Detection Kit-SPiDER-βGal (Dojindo Molecular Technologies, Inc, Rockville, MD) was used to identify senescent CD8 T cells as previously described by us^8^. Briefly, CD8 T cells were incubated 37 °C, 5% CO2 with a 1:1000 dilution of bafilomycin A-1 for 30 min before the addition of a 1:1000 dilution of SPiDER-βGal for an additional 30 min. Cells were subsequently washed with 1x PBS (Corning, Manassas, VA) and resuspended in 1xPBS + 2% FBS for surface staining. Depending on the experiment, cells were surface-stained using antibodies against CD3, CD8, CCR7, CD45RA, and PD1, subsequently washed in 1x PBS, and sorted. To isolate SA-βGal^low^ and SA-βGal^high^ CD8 T_EM_ cells, enriched CD8 T cells were sorted based on 4′,6-diamidino-2-phenylindole negative (DAPI-), (BioLegend, San Diego, CA), CD8, CCR7-, CD45RA-cells with very high (SA-βGal^high^) and low (SA-βGal^low^) signal intensities. Anti-PD1 antibodies were added for experiments analyzing features of exhaustion of CD8 T_EM_ cells. Anti-CD3 antibodies were additionally added to isolate highly pure populations of CD8 T cells for single-cell RNA sequencing experiments. All analyses were performed with FlowJo software (FlowJo 10.6.0).

### Single-cell RNA sequencing, analysis, and data mining

Relative SA-βGal low and high cells were sorted from the DAPI-, CD3+, CD8 T_N,_ T_CM,_ T_EM,_ and T_EMRA_ cells obtained from 2 healthy human donors. An approximately equal number of SA-βGal^low^ SA-βGal^high^ CD8 T cells were sorted from each differentiation state. All of the four SA-βGal^low^ differentiation states per donor were pooled into a single tube, while all of the SA-βGal^high^ differentiation states per donor were pooled into a separate tube. Gel Beads-in-Emulsion (GEMs) were generated using the NextGEM Single Cell 3’ Reagent Kits v3.1 (PN-1000128, 10x Genomics, Pleasanton, CA) following the manufacturer’s protocol. The cell/RT mix, Gel Beads, and Partitioning Oil were loaded onto a Chromium Chip G (PN-1000127) in the recommended order and processed on the Chromium Controller to generate GEMs. GEM-RT incubation, post-RT cleanup, and cDNA amplification were performed according to the user guide. Amplified cDNA was size-selected to obtain fragments suitable for 3’ Gene Expression (GEX) library preparation and Cell Surface Protein (CSP) library construction. 3’ Gene Expression and Cell Surface Protein libraries were constructed using the NextGEM Single Cell 3’ Reagent Kits v3.1 (10x Genomics) following the manufacturer’s instructions. Library concentration and quality were assessed using a Qubit 4 fluorometer with the High Sensitivity dsDNA reagent kit (Cat. No. Q33231, Thermo Fisher Scientific) and an Agilent TapeStation system with High Sensitivity DNA D1000 ScreenTape (PN 5067-5584). Final libraries were diluted and sequenced on an Illumina NovaSeq 6000 system using an S1 100-cycle flow cell (Cat. No. 20028319) with a 28/10/10/90 bp read configuration for cell barcode, sample index, and mRNA reads, respectively, as recommended by 10x Genomics. Raw FASTQ files were uploaded to the 10X Genomics cloud server and preprocessed to generate barcode, feature, and matrix files using their standard online workflow. Briefly, 10x Genomics output files were processed in R Studio using Seurat to generate Seurat files utilizing the CreateSeuratObject function. The 4 Seurat objects were subsequently merged using the merge function and preprocessed together. Cells were filtered for low quality by removing cells with <500 and >4000 features, >20000 counts, and >10% mitochondrial DNA genes; cutoffs were determined after visually inspecting the distribution of data. Following filtering, the merged dataset was log-normalized with a scale factor of 10000, and variable features were identified using the FindVariableFeatures function with an nfeature of 2000. The data were subsequently scaled using the ScaleData function in Seurat. The RunPCA function was applied to the scaled data, and the dimensions were explored using the elbowPlot function. The top 20 principal components PCs were utilized for graph-based clustering. To remove donor-specific clusters without overcorrecting, we employed Harmony with a theta of 5. Batch-corrected data, including only the top 6 PCs, were utilized to identify CD8 T cell subsets with a FindClusters resolution of 0.4. CD8 T cell clusters were annotated based on the genes used to discriminate various differentiation states in single-cell RNAseq data from human CD8 T cells as described^29^. EM clusters were isolated from the main Seurat object and processed similarly as above to generate the resulting figures. For the scRNAseq data mining, publicly available datasets were downloaded from GEO and preprocessed using the above workflow with minor modifications. Briefly, raw counts were extracted from the available data and subsetted to include only the genes found in our dataset. Datasets were then separately merged with our data, and the resulting Seurat object was down sampled to include fewer cells. The code used to preprocess and analyze the data will be made available upon publication.

### Gene expression analysis

FACS-sorted SA-βGal^low^ and SA-βGal^high^ CD8 T_EM_ cells from 6 healthy human donors were purified using the Macherery-Nagel RNA XS Plus Kit per manufacturer’s instructions (Macherery-Nagel, Duren, Germany). Equal amounts of purified RNA were used to generate cDNA using a Bio-Rad iScript cDNA synthesis kit, following the manufacturer’s instructions (Bio-Rad, Hercules, CA). Qiagen QuantiTect primers were used for CDKN1A, CDKN2A, CKDN2B, and MYC, as well as GAPDH as a reference housekeeping gene. Real-time RT-qPCR was performed on a Bio-Rad CFX96 Real-Time PCR detection system (Bio-Rad, Hercules, CA). Data were analyzed using the Bio-Rad CFX Manager software, version 3.1.

### Immunofluorescence

Immunofluorescence staining of CD8 T cells was performed as described^59^. Briefly, FACS-sorted SA-βGal^low^ and SA-βGal^high^ CD8 T_EM_ cells were collected and resuspended in 1x PBS at 10^6^ cells/ml. The cell suspension was layered on glass coverslips and was left standing for 30 min for gravity sedimentation. After 30 min, the cells were fixed in 4% formaldehyde for 15 min and permeabilized for 15 min with 0.2% Triton X-100 in PBS (PBST 0.2%). Following permeabilization, cells were incubated in blocking buffer (4% BSA in PBST 0.1%) for 1 h or overnight. Cells were subsequently incubated with primary antibodies overnight at 4 °C and then washed with PBST 0.1% (3 times, 10 minutes each). Cells were incubated with secondary antibodies for 1 h at room temperature and subsequently washed with 1x PBS (3 x 10 minutes), air-dried, and mounted on slides using DAPI-containing mounting medium (Invitrogen, Waltham, MA). Slides were imaged within 2 days of mounting.

### Telomere-ImmunoFISH

Telomere-ImmunoFISH was performed as previously described by us^60^ with slight modifications to enhance the 53BP1 signal. Briefly, cells were processed and immunostained using anti-53BP1 and secondary antibodies as described above (see “Immunofluorescence”). Following staining, cells were washed with 1x PBS (3x 10 min) and dehydrated by subsequent submersion in 70%, 90%, and 100% ethanol (3 min each). After dehydration, cells were air-dried and incubated with 0.5 μg/ml TelC-Cy3 or TelC-Cy5 PNA probes (Panagene, Korea) in hybridization buffer (70% formamide, 12 mM Tris-HCl pH = 8.0, 5 mM KCl, 1 mM MgCl2, 0.001% Triton X-100, 0.25% acetylated BSA) for 5 min at 80°C, to allow denaturation of DNA and hybridization of the telomere-specific PNA probe. After gradual cooling, cells were incubated overnight in a humidified chamber at room temperature. Subsequently, cells were washed 2 times with 70% formamide/0.6X SSC (15 min each) followed by 2 washes with 2X SSC buffer (10 min each), and 3 washes in PBST (3×10 minutes). Cells were incubated with secondary antibodies for 1h at room temperature to reinforce 53BP1 signal, washed with 1x PBS (3 x 10 minutes), air-dried, and mounted on slides as described above (See “Immunofluorescence”). Images were acquired using a Zeiss Axio Observer Z1 Inverted Phase Contrast Fluorescence Microscope and a 63X or 40X oil-immersive lens. Images were obtained using Z-stacks and analyzed with Zeiss ZEN 2.5 (blue edition) software. For each experiment, at least 100 cells were analyzed to quantify the number of 53BP1 foci per cell, as well as co-localizations of 53BP1 foci with telomeres. Only distinct, well-formed 53BP1 foci in each plain focus of the Z-stack were counted. T cells with granular, non-focal 53BP1 staining pattern were excluded from the analyses.

### Antibodies

PerCP/Cy5.5 anti-CD3 (HIT3a), PE anti-CD8a (RPA-T8), APC anti-CD54RA (HI100), BV711 anti-CD54RA (HI100,) PE Cy7 anti-CCR7 (G043H7), APC Cy7 anti-CCR7 (G043H7), AF700 anti-Perforin (B-D48), APC anti-Granzyme A (CB9), APC/Fire 750 anti-CCR9 (LO53E8), BV510 anti CX3C31 (2A9-1), BV605 anti-IFNy (B27), BV650 anti-CXCR3 (G025H7), BV750 anti-TNF (Mab11), BV785 anti-IL2 (MQ1-17H12), PacBlue anti-CXCR2 (5E8), PE anti-TCF7 (7F11A10), PE-Dazzle 594 anti-Granzyme B (QA16A02), PE-Fire 700 anti-CD45RA (HI100), PE-Fire 810 anti-PD1 (A17188B), PerCP-Fire 780 anti-CD28 (S20013B), Spark Blue 574 anti-CXCR1 (8F1), Spark Plus B550 anti-LAMP1 (H4A3), Spark Plus UV395 anti-CCR6 (G034E3), PE anti-PD-L1 (29E.2A3) were purchased from BioLegend (San Diego, CA) BUV805 anti-CD8 (SK1) and BUV737 anti-PD1 (EH12.1) were purchased from BD Biosciences (Franklin Lakes, NJ). PerCP-eFluor 710 anti Granzyme K(G3H69) was purchased from Invitrogen (Waltham, MA). For immunofluorescence experiments: anti-p15 (D-12, Santa Cruz, Dallas, TX), anti-p16 (JC8, GeneTex, San Antonio, TX), anti-53BP1 (polyclonal; Novus, Littleton, CO), anti-macroH2A (in house, a gift from the Peter Adams lab). Secondary antibodies used for immunofluorescence staining were as follows: Cy3 donkey anti-mouse (Jackson ImmunoResearch, West Grove, PE) and Alexa Fluor 488 goat anti-rabbit (Invitrogen, Waltham, MA).

### Senescence-Associated βGalactosidase cell staining (X-Gal)

FACS-sorted SA-βGal_low_ and SA-βGal^high^ CD8 T_EM_ cells were gravity sedimented as described above (see “Immunofluorescence”) and subsequently processed using the senescence β-Galactosidase staining kit from Cell Signaling Technologies (Danvers, MA, USA) per manufacturer’s protocol. Briefly, cells were fixed in the provided 1x fixative solution for 10 minutes, then washed twice in 1x PBS and incubated overnight at 37 °C in a dry incubator (no CO_2_) with 1 mL of the provided β-Galactosidase staining solution at a final pH of 6. Coverslips containing cells were monitored for the development of a blue color, and reactions were stopped by removing the staining solution, washing twice with 1x PBS, and preserving the cells on a glass slide with glycerol.

### *In vitro* proliferation

Proliferation of FACS-sorted SA-βGal^low^ and SA-βGal^high^ CD8 T_EM_ cells was monitored via CellTrace Violet (CTV) (ThermoFisher, Waltham, MA). CTV staining was performed according to the manufacturer’s protocol immediately following FACS sort. Labeled cells were stimulated using anti-CD3/CD28 dynabeads (ThermoFisher, Waltham, MA) (1 bead : 1 cell ratio) or stimulated with anti-CD3 dynabeads (ThermoFisher, Waltham, MA) (1 bead : 1 cell ratio) supplemented with either 25ng/ml of IL2, IL7, 15 or all 3 (StemCell Technologies, Cambridge, MA) for 7 days at 37°C in a humidified atmosphere with 5% CO_2_. All T cells were grown in RPMI-1640 containing 10% FBS and 1x penicillin streptomycin. After 7 days, cells were collected, stained with DAPI, and analyzed on a BD Symphony (BD Biosciences, Franklin Lakes, NJ).

### Bulk RNA sequencing and *ex vivo* stimulation

FACS-sorted SA-βGal^low^ and SA-βGal^high^ CD8 T_EM_ cells from 6 healthy human donors were either collected for RNA extraction or stimulated with anti-CD3/CD28 (1 bead:1 cell ratio) dynabeads (ThermoFisher, Waltham, MA) for an additional 16 hours before being collected for RNA extraction. RNA was purified using a Macherey-Nagel RNA XS Plus kit according to the manufacturer’s instructions (Macherey-Nagel, Duren, Germany). RNA integrity was evaluated in a Bioanalyzer 2100 system, and only RNA with an integrity number of >= 7 was used for library preparation. Libraries were constructed using the SMARTer Stranded V2 according to the manufacturer’s instructions (TakaraBio, Mountain View, CA). Paired-end sequencing was performed on an Illumina HiSeq 2500 instrument. At least 40 million reads per sample (20 million per strand) were obtained and used for downstream analyses.

### Preprocessing of high-throughput bulk RNA sequencing data

Paired-end RNA-seq reads were processed as previously described by us^61^ and aligned to the GRCh38.d1.v1 version of the human genome using bowtie2^62^ using the local mode (RNA-seq). Low-quality reads and adapters were removed using fastq-mcf v.1.0.5 and cutadapt. Alignments were further processed using samtools v.1.1.1.PCR duplicates and optical duplicates were removed with PicardTools v.2.2.2. Reads were counted using summarized overlaps and normalized with DESeq2 for visualization.

### Differential expression

Differentially expressed genes (DEGs) were identified from Bulk RNA-seq data, using DESeq2. Raw reads were internally normalized by DESeq2 using the median of ratios method. Reads per exon were obtained using the GRCh38.107 genome model and quantified using the *summarizeOverlaps* package. Peaks/genes with at least 10 reads in at least 4 libraries per phenotypic state were kept. Correction of batch effects was performed with limma ^63^ using the source of samples (batch, freshly isolated or isolated after ∼16 hours) as the surrogate variable. Data transformation (regularized-log [rld] transformation), exploratory visualization (PCA and hierarchical clustering), and differential expression analysis were performed with DESeq2 using the default parameters as previously described and base R functions. Only highly significant DEGs using an adjusted *p*-value filter of 0.1 or 0.05 were considered.

### Functional over representation analysis

The DEGs identified between the scRNAseq T_EM_ clusters and WGCNA modules were characterized by over-representation tests using the clusterprofiler package. A collection of pathways from Hallmark, KEGG, Reactome, GO, BioCarta and Wikipathways was used as input. Statistical significance was calculated by a hypergeometric test with a cut-off of an adjusted *p*-value of <= 0.1 with Benjamini–Hochberg correction.

### Self-organizing maps (SOMs)

SOM expression portraits were generated using the unsupervised machine learning method deployed in oposSOM ^41^. Metagenes were visualized in a 60 x 60 grid of rectangular topology, wherein expression portraits are projected by metagene distance matrix similarity using a logarithmic fold-change scale.

### WGCNA on differentially expressed genes

Differentially expressed genes (DEGs) identified with DESeq2 were used as input for unsupervised clustering using WGCNA. We used the “signed” option with default parameters, except for the soft thresholding power for RNA-seq data, which was set to 17. The minimum size of DEGs for the initial set of modules was set to either 50, 200 or 600 depending on subset of genes. RNA-seq datasets were merged by a dissimilarity threshold of 0.5 or 0.2 depending on subset of genes. WGCNA modules were functionally profiled with clusterProfiler using the Molecular Signatures Database Hallmark gene sets, GO, and Reactome pathways. Statistical significance was calculated by a hypergeometric test with a cut-off of an adjusted *p*-value of <= 0.1 with Benjamini–Hochberg correction.

### *Ex vivo* stimulation, surface staining, intracellular staining, and cyCONDOR analysis

For *ex vivo* stimulation, FACS-sorted SA-βGal^low^ and SA-βGal^high^ CD8 T_EM_ cells from 6 healthy human donors were divided into either unstimulated, anti-CD3/CD28 bead-stimulated (1cell :1 bead ratio), or PMA (500ng) plus Ionomycin (1ug) for 5 hours in the presence of Golgi-block (monensin, STEMCELL Technologies, Vancouver, BC) and anti-CD107a. Following the 5-hour stimulation, cells were washed in blocking buffer (1xPBS +2% FBS) and surface-stained for IL7R, CXCR3, CD28, CXCR1, PD1, CCR6, CCR9, CD16, CX3CR1, and CXCR2 for 30 minutes at 37 °C while protected from light. Cells were subsequently washed in 1x PBS and stained with LIVE/DEAD Fixable Blue Dead Cell Stain (ThermoFisher, Waltham, MA) per manufacturer’s instructions, after which the cells were washed in 1X PBS and fixed using True-Nuclear Transcription Factor Buffer Set (BioLegend, San Diego, CA) per manufacturer’s protocol. Cells were left in 1x fixing solution overnight, washed in the provided 1x permeabilization solution (2x 10minutes), and immunostained in the same solution using antibodies against GrA, GrB, GrK, Perforin, TCF7, IFNy, TNF and Il2 for 30 minutes at room temperature while protected from light. Cells were subsequently washed in blocking buffer (1xPBS +2% FBS, 2 x10 minutes) and immediately analyzed on a BD Symphony (BD Biosciences, Franklin Lakes, NJ). For CyCONDOR analysis, freshly isolated and live CD8 T cell populations, sorted using a BD Symphony from 6 donors, were exported from FlowJo according to the developers’ recommendations. 2,500 cells from each donor were randomly extracted, auto-logically transformed, and used for downstream analysis. Batch effects were corrected for using the integrated harmony package, with batch being defined as different days on which samples were stained and acquired. UMAP projections, stacked bar graphs, scaled marker expressions, and box plots were plotted using standard cyCONDOR workflows.

### Cell culture

GM21 fibroblasts were obtained from the Coriell Institute and cultured in RPMI-1640 containing 10% FBS and 1x penicillin/streptomycin at 37 °C and 2% oxygen. For DNA damage-induced senescence, GM21 fibroblasts were treated with a single dose of 20 μM etoposide for 2 days, followed by culturing for at least 10 days in the absence of drug before use. Media was changed every 2-3 days. Raji B cell lymphoma cells were a gift from the Dongfang Liu laboratory (NJMS-Rutgers) and cultured in RPMI-1640 containing 10% FBS and 1x penicillin streptomycin at 37 °C, 21% oxygen. Raji cells were passaged every 2-3 days in fresh media.

### Cytotoxicity assays and analysis

Senescent GM21 fibroblasts were trypzized using 0.05% trypsin (Corning, Manassas, VA) and seeded in xCELLigence 96-well E-plates (Agilent Technologies, CA, USA) using a multichannel pipette to minimize variability between wells, 3,000 cells/well. E-plates were loaded into the xCELLigence system, which was kept in a 37 °C, 5% CO_2_ incubator. Cells were allowed to adhere for 24 hours, with measurements taken every minute. After 24h unstimulated or 3-day soluble anti-CD3/CD28 stimulated FACS-sorted SA-βGal^low^ and SA-βGal^high^ CD8 T_EM_ cells were added to each well at effector to target ratios as indicated in figure legends; for all experiments involving senescent fibroblasts, E:T is 25:1. Cytolysis was monitored over a 24-hour period. The technical efficiency of the instrument was evaluated by ensuring full lysis of target cells in 0.1% Triton-X (not shown). All experimental conditions were conducted in duplicates or triplicates wherever possible. Raw cell index values were exported as CSV files and analyzed in Excel (Microsoft, WA, USA). Briefly, the raw cell index values of each well were normalized to the raw cell index values of that same well immediately before adding effector cells to obtain the normalized cell index (NCI). NCI values were averaged by across technical replicates. To obtain relative cytolysis, the following formula was used: [NCI(control) – NCI(treatment)]/NCI(control) x 100. Negative relative cytolysis values, reflecting senescent fibroblast movement and increased adherence, were set to 0; for anti-PD1 experiments smallest NCI values were subtracted from each time point. For PD-1 blockade experiments, cells were stimulated for 3 days using soluble anti-CD3/CD28 antibodies and subsequently incubated for 30 minutes with anti-PD-1 antibodies before being added to senescent fibroblasts. For flow cytometry-based cytotoxicity, Raji cells were stained with Calcein AM Blue according to the manufacturer’s protocol and subsequently cultured with pre-stimulated (as above) FACS-sorted SA-βGal^low^ and SA-βGal^high^ CD8 T_EM_ cells for 16 hours at various effector-to-target ratios, as indicated in the figure legend. All conditions were performed in duplicate wherever possible. Counting beads were utilized to calculate the absolute number of cells, after which the % cytolysis was calculated with the following formula: absolute cell number (control) – absolute cell number (treatment)]/ absolute cell number (control) x 100.

### GSEA

GSEA of unstimulated SA-βGal^low^ and SA-βGal^high^ CD8 T_EM_ cells signatures derived from our transcriptomic dataset was performed on transcriptomes of peripheral T_EM_ cells from: younger and older individuals (GEO: GSE179613)^44^, individuals with rheumatoid arthritis (GEO: GSE118829)^46^, individuals with stage IV melanoma (GEO: GSE104744)^47^, or from the peripheral CD8 T cells of smokers (GEO: GSE118829)^45^. We also utilized the gene signature from single-cell RNA sequencing data of T cells derived from nonresponders to anti-PD1 therapy^50^ and applied it to our ranked unstimulated SA-βGal^low^ and SA-βGal^high^ CD8 T_EM_ cell transcriptomes. All analyses were conducted using the GSEA GUI version 4.4.0.

## Quantification and statistical analysis

Except for sequencing analyses, all statistical tests used in this study were performed in GraphPad Prism, as indicated in the legend of their respective figures. Unless otherwise stated, all experiments were performed with at least 3 biological replicates (at least 3 independent donors) and confirmed with at least 2 independent experiments.

## Supplemental Figures

**Figure S1.**
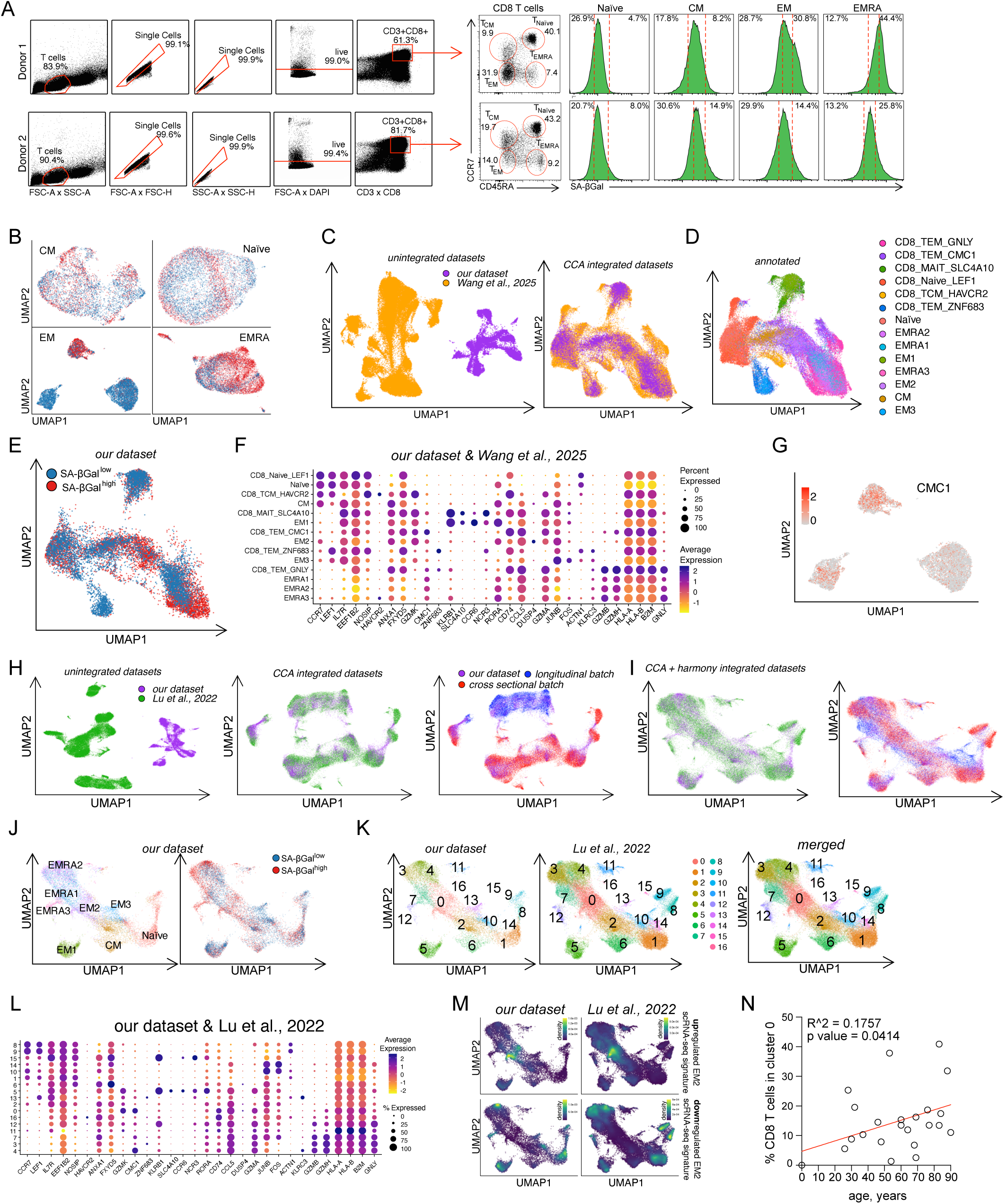
**A**. Representative FACS scatter plots demonstrating the gating strategy to identify SA-βGal^low^ and SA-βGal^high^ CD3+CD8 T cells in four differentiation states, as defined by CD45RA and CCR7 expression. T_Naive_: CD45RA^+^/CCR7^+^; T_CM_: CD45RA^-^/CCR7^+^; T_EM_: CD45RA^-^/CCR7^-^; T_EMRA_: CD45RA^+^/CCR7^-^. **B**. UMAP Projections of subsetted CD8 T_Naive_, T_CM_, T_EM_, and T_EMRA._ with cells colored based on the SA-βGal sorting gate (blue: SA-βGal^low^; red: SA-βGal^high^). **C**. UMAP projection of CD3+/CD8 T cells derived from our dataset (purple) and from *Wang et al., 2025*^38^ (gold) before (left) and after (right) batch correction. **D**. CCA batch corrected data projected on UMAP embedding and annotated with our, or Wang et al., 2025 cell annotation. **E**. Same UMAP as in (A), showing cells only from our dataset and colored based on the SA-βGal sorting gate (blue: SA-βGal^low^; red: SA-βGal^high^). **F**. Dotplot of CD3+/CD8 T cell clusters identified in our study and the Wang et al., 2025 study. Average expression levels and percentage of cells expressing the indicated marker genes are shown. **G**. UMAP projection of CD8 T_EM_ cell clusters showing gene expression of CMC1. **H**. UMAP projection of CD3+/CD8 T cells derived from our dataset (purple) and that from Lu et al., 2022 (green) before (left) and after (middle & right) CCA batch correction. **I**. UMAP projection of CD3+/CD8 T cells derived from our dataset (purple) and that from Lu et al., 2022 (green) after additional harmony batch correction. **J**. Same UMAP as in (I), showing cells only from our dataset and colored based on our cluster annotation (left) and on the SA-βGal sorting gate (right; blue: SA-βGal^low^; red: SA-βGal^high^). **K**. Same UMAP as in (I) annotated by new clusters and split by our dataset (left) Lu et. al. 2022 (center) or merged (right). **L**. Dotplot of CD3+/CD8 T cell clusters identified in our and the Lu et al. 2022 study. Average expression levels and percentage of cells expressing the indicated marker genes are shown. **M**. GSEA using AUCell scoring, transformed into density scores, mapped onto UMAP projection for EM2-specific upregulated (top row) or downregulated (bottom row) genes in our dataset (left column) or that from *Lu et al., 2025* (right). **O**. Quantification of the percentage of cells in cluster 0 out of the total CD8 T cell population in healthy human donors at indicated ages.

**Figure S2.**
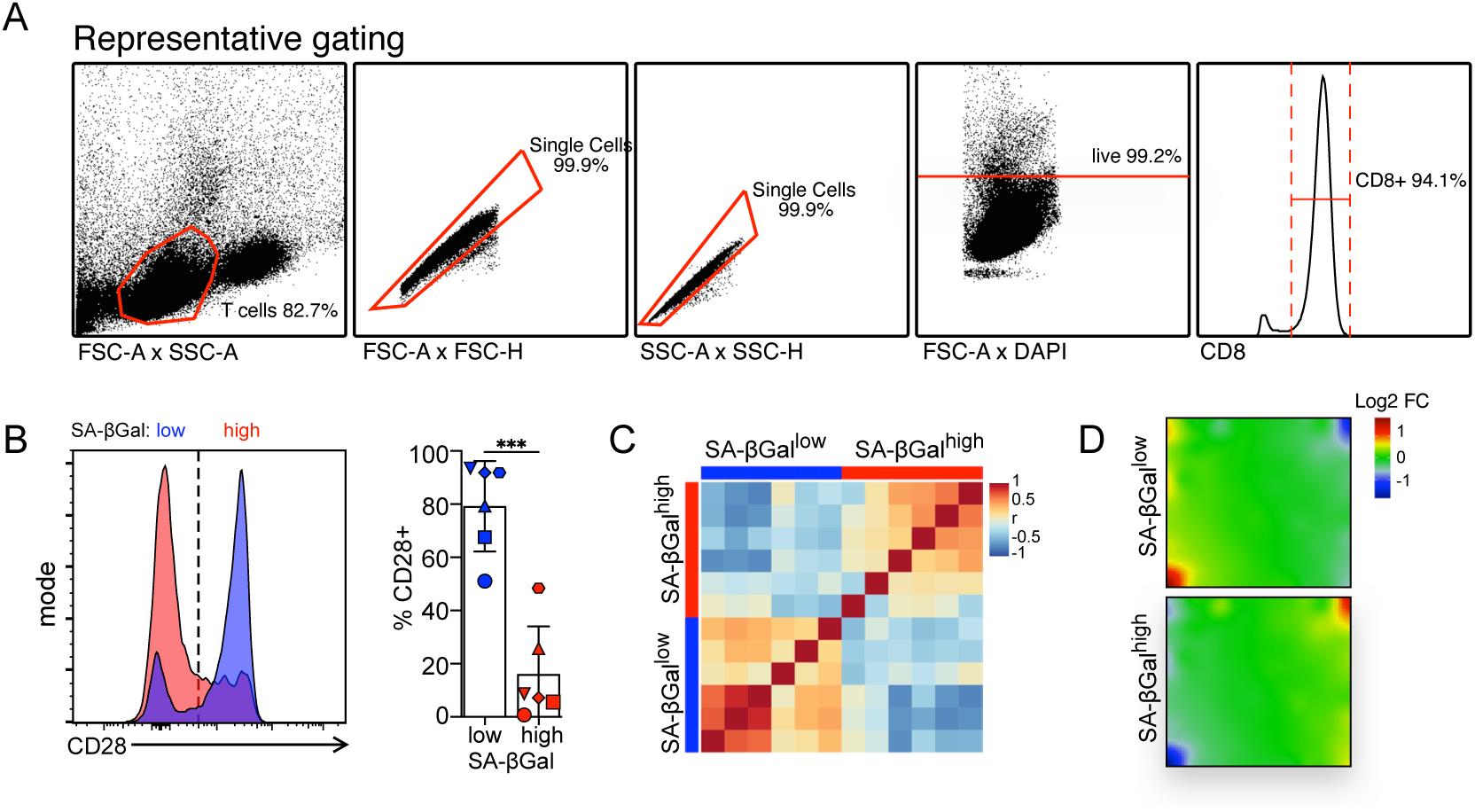
**A**. Representative FACS scatter plots demonstrating the gating strategy to sort and isolate CD8 T cells from peripheral blood mononuclear cells. **B**. Representative FACS histogram showing CD28 expression in SA-βGal^low^ (blue) and SA-βGal^high^ (red) T_EM_ cells. Bar graph show quantification of the fraction of CD28 expressing cells in indicated cell populations (mean +/- SD). Shapes of data points in the bar graphs correspond to unique donors. \*\*\**p* ≤0.001 by paired two-tailed t test. **C**. Heatmap showing the correlation (r) between the transcriptomes of SA-βGal^low^ and SA-βGal^high^ CD8 T_EM_ cells, as indicated, from 6 donors. **D**. Averaged self-organizing maps (SOMs) of transcriptomes from freshly isolated unstimulated SA-βGal^low^ and SA-βGal^high^ T_EM_ cells.

**Figure S3.**
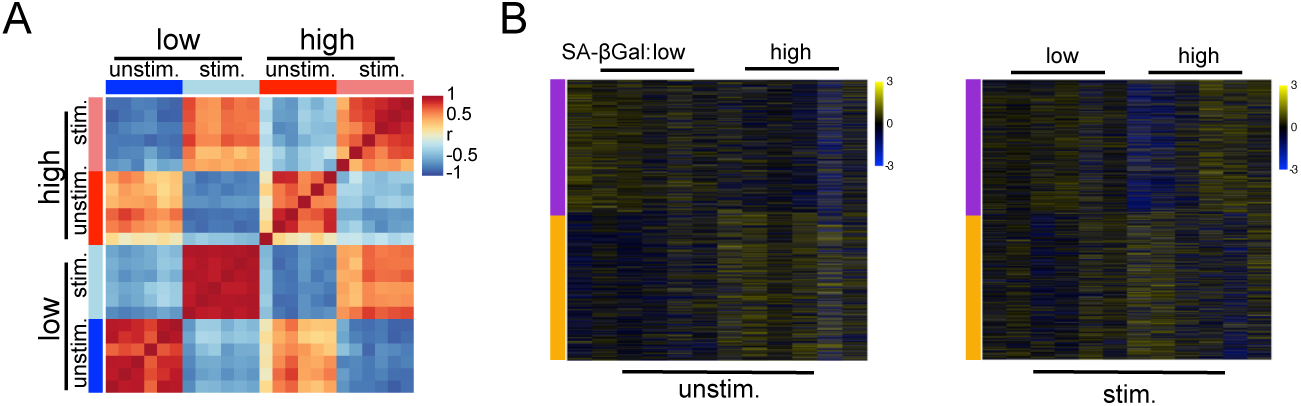
**A**. Heatmap showing the correlation (r) between the transcriptomes of SA-βGal^low^ and SA-βGal^high^ CD8 T_EM_ cells, stimulated with anti-CD3/CD28 antibodies or not as indicated, from 6 healthy human donors. **B**. WGCNA heatmaps of indicated color-coded modules showing only the shared, stimulation-specific DEGs.

**Figure S4.**
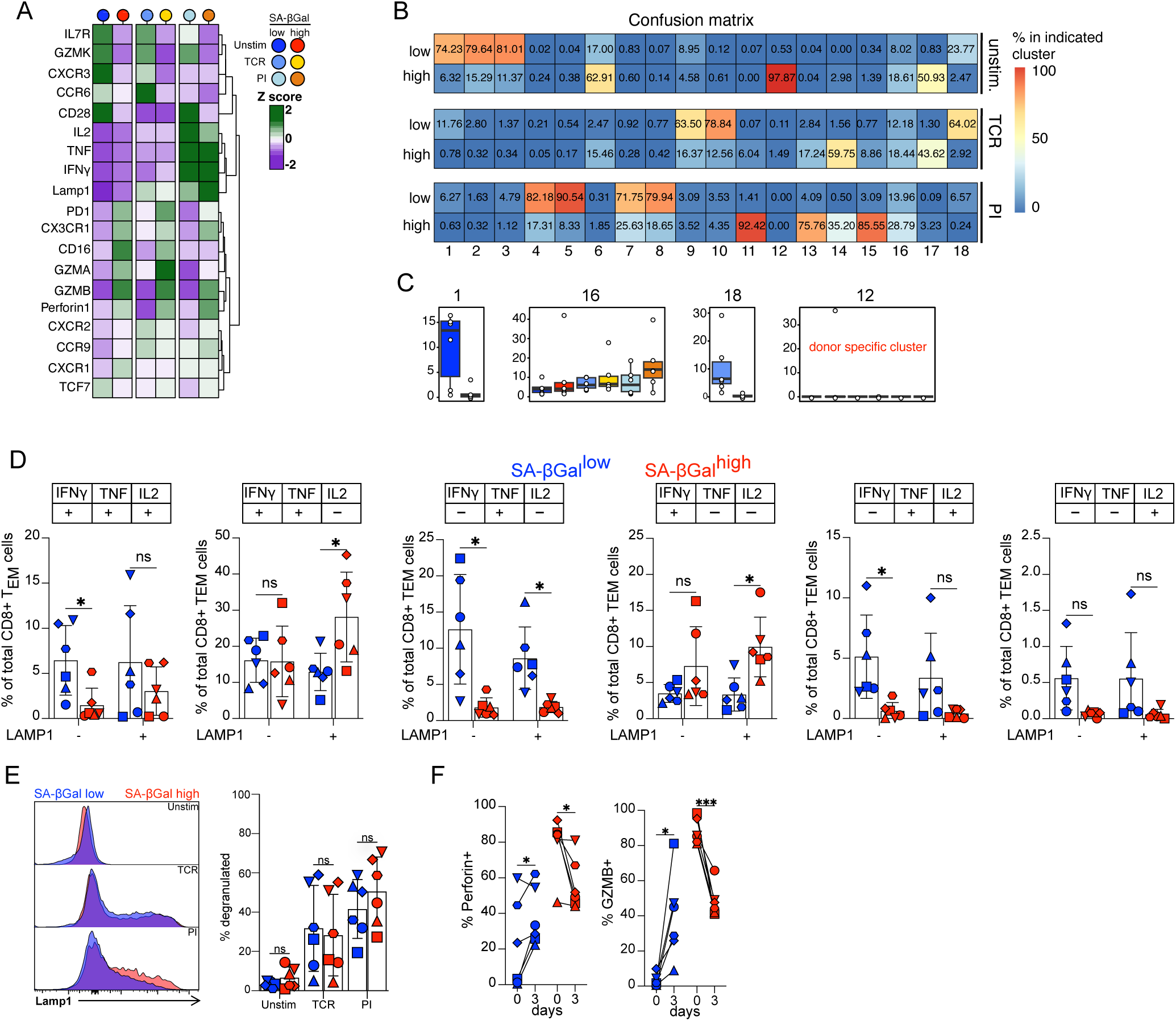
**A**. Expression pattern heatmaps (Z score) of indicated 19 markers of SA-βGal^low^ (blue) and SA-βGal^high^ (red) CD8 T_EM_ cells, that were unstimulated (unsim.) and stimulated with anti-CD3/CD28 antibodies (TCR) or PMA/Ionomycin (PI) as indicated. **B**. Confusion matrix demonstrating the percent contribution to a given FlowSOM cluster (numbered columns) from SA-βGal^low^ (low) and SA-βGal^high^ (high) CD8 T_EM_ cells that were unstimulated (unsim.) and stimulated with anti-CD3/CD28 antibodies (TCR) or PMA/Ionomycin (PI) as indicated. **C**. Box plots showing the % of sorted SA-βGal^low^ and SA-βGal^high^ CD8 T_EM_ cells within select FlowSOM clusters. Colors of boxes represent different stimulation conditions, as in Figure 4A. Each circle represents a unique donor. **D**. Quantification of the indicated cell populations derived from Boolean gating in SA-βGal^low^ (blue) and SA-βGal^high^ (red) T_EM_ cells stimulated with PMA/Ionomycin for 5 h. Bar graph is represented as mean +/- SD; Shapes of data points in the bar graphs correspond to unique donors. *\*adj.p* ≤0.05 by paired two-tailed t test with Holm-Šídák correction. **E**. Representative FACS histogram (left) and quantification (right) showing externalized CD107a in FACS-sorted SA-βGal^low^ and SA-βGal^high^ T_EM_ cells that were either unstimulated (unstim.), or stimulated for 5 h with anti-CD3/CD28 antibodies (TCR) or PMA/Ionomycin (PI). n = 6. Bar graphs are represented as mean +/- SD. Shapes of data points in the bar graphs correspond to unique donors. **F**. Quantification of Perforin and GzmB expression in in FACS sorted SA-βGal^low^ and SA-βGal^high^ T_EM_ cells from 6 donors on days 0 and 3. Shapes of data points in the bar graphs correspond to unique donors. *\*Adj.p* ≤0.05, ^\*\**\**^*adj.p* ≤0.001 by two-tailed paired t test.

**Figure S5.**
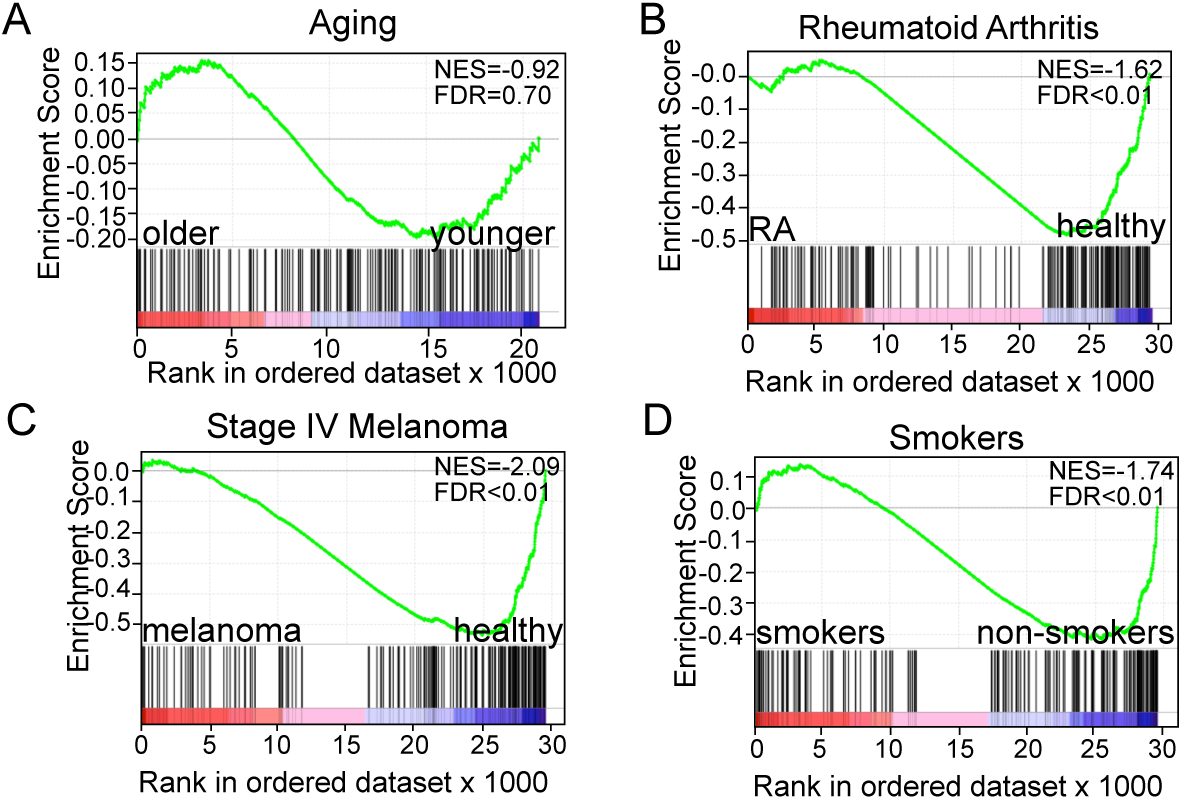
Gene set enrichment analyses (GSEA) showing normalized enrichment score (NES) plots and false discovery rate (FDR) values for our non-senescent CD8 T_EM_ cell gene signature that enriches in normalized and ranked transcriptiomes of peripheral blood CD8 T_EM_ cells isolated from younger healthy human donors (**A**), healthy controls in groups (**B-C**), and from peripheral blood CD8 T cells of non-smokers (**D**).

**Figure S6.**
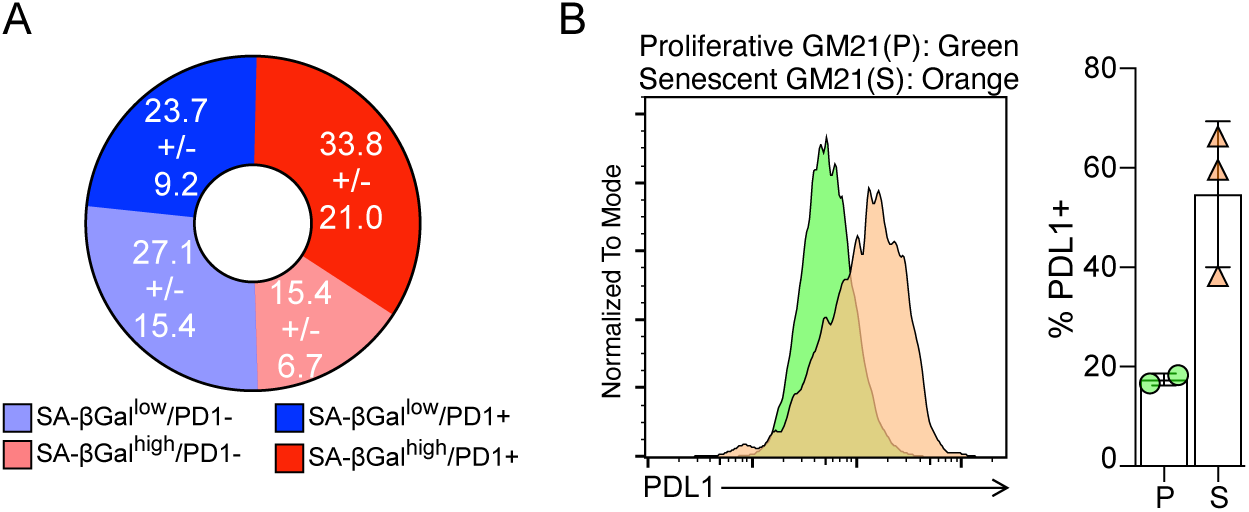
**A**. Pie chart showing the frequency of SA-βGal^low^/PD1- (light blue), SA-βGal^low^/PD1+ (dark blue), SA-βGal^high^/PD1-(light red), and SA-βGal^high^, PD1+ (dark red) CD8 T_EM_ cells. Mean +/- SD is indicated in respective segments **B**. Representative FACS histogram (left) and quantification (right) of PDL1 expression in non-senescent (green) or senescent (orange) GM21 humasn dermal fibroblasts. Bar graph is represented as mean +/- SD.

## Notes

### Competing Interest Statement

The authors have declared no competing interest.

## References

1. Di Micco, R., Krizhanovsky, V., Baker, D., and d’Adda di Fagagna, F. (2021). Cellular senescence in ageing: from mechanisms to therapeutic opportunities. Nat Rev Mol Cell Biol 22, 75–95. 10.1038/s41580-020-00314-w.

2. Prata, L., Ovsyannikova, I.G., Tchkonia, T., and Kirkland, J.L. (2019). Senescent cell clearance by the immune system: Emerging therapeutic opportunities. Semin Immunol 40, 101275. 10.1016/j.smim.2019.04.003.

3. Munoz-Espin, D., and Serrano, M. (2014). Cellular senescence: from physiology to pathology. Nat Rev Mol Cell Biol 15, 482–496. 10.1038/nrm3823.

4. Liu, Z., Liang, Q., Ren, Y., Guo, C., Ge, X., Wang, L., Cheng, Q., Luo, P., Zhang, Y., and Han, X. (2023). Immunosenescence: molecular mechanisms and diseases. Signal Transduction and Targeted Therapy 8, 200. 10.1038/s41392-023-01451-2.

5. Ovadya, Y., Landsberger, T., Leins, H., Vadai, E., Gal, H., Biran, A., Yosef, R., Sagiv, A., Agrawal, A., Shapira, A., et al. (2018). Impaired immune surveillance accelerates accumulation of senescent cells and aging. Nat Commun 9, 5435. 10.1038/s41467-018-07825-3.

6. Desdin-Mico, G., Soto-Heredero, G., Aranda, J.F., Oller, J., Carrasco, E., Gabande-Rodriguez, E., Blanco, E.M., Alfranca, A., Cusso, L., Desco, M., et al. (2020). T cells with dysfunctional mitochondria induce multimorbidity and premature senescence. Science 368, 1371–1376. 10.1126/science.aax0860.

7. Yousefzadeh, M.J., Flores, R.R., Zhu, Y., Schmiechen, Z.C., Brooks, R.W., Trussoni, C.E., Cui, Y., Angelini, L., Lee, K.A., McGowan, S.J., et al. (2021). An aged immune system drives senescence and ageing of solid organs. Nature 594, 100–105. 10.1038/s41586-021-03547-7.

8. Martinez-Zamudio, R.I., Dewald, H.K., Vasilopoulos, T., Gittens-Williams, L., Fitzgerald-Bocarsly, P., and Herbig, U. (2021). Senescence-associated beta-galactosidase reveals the abundance of senescent CD8 T cells in aging humans. Aging Cell 20, e13344. 10.1111/acel.13344.

9. Pereira, B.I., Devine, O.P., Vukmanovic-Stejic, M., Chambers, E.S., Subramanian, P., Patel, N., Virasami, A., Sebire, N.J., Kinsler, V., Valdovinos, A., et al. (2019). Senescent cells evade immune clearance via HLA-E-mediated NK and CD8(+) T cell inhibition. Nat Commun 10, 2387. 10.1038/s41467-019-10335-5.

10. Munoz, D.P., Yannone, S.M., Daemen, A., Sun, Y., Vakar-Lopez, F., Kawahara, M., Freund, A.M., Rodier, F., Wu, J.D., Desprez, P.Y., et al. (2019). Targetable mechanisms driving immunoevasion of persistent senescent cells link chemotherapy-resistant cancer to aging. JCI Insight 5. 10.1172/jci.insight.124716.

11. Sturmlechner, I., Zhang, C., Sine, C.C., van Deursen, E.J., Jeganathan, K.B., Hamada, N., Grasic, J., Friedman, D., Stutchman, J.T., Can, I., et al. (2021). p21 produces a bioactive secretome that places stressed cells under immunosurveillance. Science 374, eabb3420. 10.1126/science.abb3420.

12. Wang, T.W., Johmura, Y., Suzuki, N., Omori, S., Migita, T., Yamaguchi, K., Hatakeyama, S., Yamazaki, S., Shimizu, E., Imoto, S., et al. (2022). Blocking PD-L1-PD-1 improves senescence surveillance and ageing phenotypes. Nature. 10.1038/s41586-022-05388-4.

13. Marin, I., Boix, O., Garcia-Garijo, A., Sirois, I., Caballe, A., Zarzuela, E., Ruano, I., Stephan-Otto Attolini, C., Prats, N., Lopez-Dominguez, J.A., et al. (2022). Cellular senescence is immunogenic and promotes anti-tumor immunity. Cancer Discov. 10.1158/2159-8290.CD-22-0523.

14. Appay, V., van Lier, R.A., Sallusto, F., and Roederer, M. (2008). Phenotype and function of human T lymphocyte subsets: consensus and issues. Cytometry A 73, 975–983. 10.1002/cyto.a.20643.

15. Pereira, B.I., De Maeyer, R.P.H., Covre, L.P., Nehar-Belaid, D., Lanna, A., Ward, S., Marches, R., Chambers, E.S., Gomes, D.C.O., Riddell, N.E., et al. (2020). Sestrins induce natural killer function in senescent-like CD8(+) T cells. Nat Immunol 21, 684–694. 10.1038/s41590-020-0643-3.

16. Laphanuwat, P., Gomes, D.C.O., and Akbar, A.N. (2023). Senescent T cells: Beneficial and detrimental roles. Immunol Rev 316, 160–175. 10.1111/imr.13206.

17. Chou, J.P., and Ejros, R.B. (2013). T cell replicative senescence in human aging. Curr Pharm Des 19, 1680–1698.

18. Slaets, H., Veeningen, N., de Keizer, P.L.J., Hellings, N., and Hendrix, S. (2024). Are immunosenescent T cells really senescent? Aging Cell 23, e14300. 10.1111/acel.14300.

19. Baessler, A., and Vignali, D.A.A. (2024). T Cell Exhaustion. Annu Rev Immunol 42, 179–206. 10.1146/annurev-immunol-090222-110914.

20. Giles, J.R., Globig, A.M., Kaech, S.M., and Wherry, E.J. (2023). CD8(+) T cells in the cancer-immunity cycle. Immunity 56, 2231–2253. 10.1016/j.immuni.2023.09.005.

21. Suram, A., and Herbig, U. (2014). The replicometer is broken: telomeres activate cellular senescence in response to genotoxic stresses. Aging Cell 13, 780–786. 10.1111/acel.12246.

22. Hewitt, G., Jurk, D., Marques, F.D., Correia-Melo, C., Hardy, T., Gackowska, A., Anderson, R., Taschuk, M., Mann, J., and Passos, J.F. (2012). Telomeres are favoured targets of a persistent DNA damage response in ageing and stress-induced senescence. Nat Commun 3, 708. 10.1038/ncomms1708.

23. Fumagalli, M., Rossiello, F., Clerici, M., Barozzi, S., Cittaro, D., Kaplunov, J.M., Bucci, G., Dobreva, M., Matti, V., Beausejour, C.M., et al. (2012). Telomeric DNA damage is irreparable and causes persistent DNA-damage-response activation. Nature cell biology 14, 355–365.

24. Hernandez-Segura, A., Nehme, J., and Demaria, M. (2018). Hallmarks of Cellular Senescence. Trends Cell Biol 28, 436–453. 10.1016/j.tcb.2018.02.001.

25. Lee, B.Y., Han, J.A., Im, J.S., Morrone, A., Johung, K., Goodwin, E.C., Kleijer, W.J., DiMaio, D., and Hwang, E.S. (2006). Senescence-associated beta-galactosidase is lysosomal beta-galactosidase. Aging Cell 5, 187–195.

26. Gorgoulis, V., Adams, P.D., Alimonti, A., Bennett, D.C., Bischof, O., Bishop, C., Campisi, J., Collado, M., Evangelou, K., Ferbeyre, G., et al. (2019). Cellular Senescence: Defining a Path Forward. Cell 179, 813–827. 10.1016/j.cell.2019.10.005.

27. Turano, P.S., Akbulut, E., Dewald, H.K., Vasilopoulos, T., Fitzgerald-Bocarsly, P., Herbig, U., and Martinez-Zamudio, R.I. (2025). Epigenetic mechanisms regulating CD8 T cell senescence in aging humans. bioRxiv. 10.1101/2025.01.17.633634.

28. Sallusto, F., Lenig, D., Forster, R., Lipp, M., and Lanzavecchia, A. (1999). Two subsets of memory T lymphocytes with distinct homing potentials and ejector functions. Nature 401, 708–712. 10.1038/44385.

29. Lu, J., Ahmad, R., Nguyen, T., Cifello, J., Hemani, H., Li, J., Chen, J., Li, S., Wang, J., Achour, A., et al. (2022). Heterogeneity and transcriptome changes of human CD8(+) T cells across nine decades of life. Nat Commun 13, 5128. 10.1038/s41467-022-32869-x.

30. Billerbeck, E., Kang, Y.H., Walker, L., Lockstone, H., Grafmueller, S., Fleming, V., Flint, J., Willberg, C.B., Bengsch, B., Seigel, B., et al. (2010). Analysis of CD161 expression on human CD8 T cells defines a distinct functional subset with tissue-homing properties. Proceedings of the National Academy of Sciences of the United States of America 107, 3006–3011. 10.1073/pnas.0914839107.

31. Konduri, V., Oyewole-Said, D., Vazquez-Perez, J., Weldon, S.A., Halpert, M.M., Levitt, J.M., and Decker, W.K. (2020). CD8(+)CD161(+) T-Cells: Cytotoxic Memory Cells With High Therapeutic Potential. Front Immunol 11, 613204. 10.3389/fimmu.2020.613204.

32. van Wilgenburg, B., Loh, L., Chen, Z., Pediongco, T.J., Wang, H., Shi, M., Zhao, Z., Koutsakos, M., Nussing, S., Sant, S., et al. (2018). MAIT cells contribute to protection against lethal influenza infection in vivo. Nat Commun 9, 4706. 10.1038/s41467-018-07207-9.

33. Zhou, X., and Xue, H.H. (2012). Cutting edge: generation of memory precursors and functional memory CD8 T cells depends on T cell factor-1 and lymphoid enhancer-binding factor-1. J Immunol 189, 2722–2726. 10.4049/jimmunol.1201150.

34. Hu, G., and Chen, J. (2013). A genome-wide regulatory network identifies key transcription factors for memory CD8(+) T-cell development. Nat Commun 4, 2830. 10.1038/ncomms3830.

35. Dominguez, C.X., Amezquita, R.A., Guan, T., Marshall, H.D., Joshi, N.S., Kleinstein, S.H., and Kaech, S.M. (2015). The transcription factors ZEB2 and T-bet cooperate to program cytotoxic T cell terminal dijerentiation in response to LCMV viral infection. J Exp Med 212, 2041–2056. 10.1084/jem.20150186.

36. Omilusik, K.D., Best, J.A., Yu, B., Goossens, S., Weidemann, A., Nguyen, J.V., Seuntjens, E., Stryjewska, A., Zweier, C., Roychoudhuri, R., et al. (2015). Transcriptional repressor ZEB2 promotes terminal dijerentiation of CD8 ejector and memory T cell populations during infection. J Exp Med 212, 2027–2039. 10.1084/jem.20150194.

37. Bignon, A., Regent, A., Klipfel, L., Desnoyer, A., de la Grange, P., Martinez, V., Lortholary, O., Dalloul, A., Mouthon, L., and Balabanian, K. (2015). DUSP4-mediated accelerated T-cell senescence in idiopathic CD4 lymphopenia. Blood 125, 2507–2518. 10.1182/blood-2014-08-598565.

38. Wang, Y., Li, R., Tong, R., Chen, T., Sun, M., Luo, L., Li, Z., Chen, Y., Zhao, Y., Zhang, C., et al. (2025). Integrating single-cell RNA and T cell/B cell receptor sequencing with mass cytometry reveals dynamic trajectories of human peripheral immune cells from birth to old age. Nat Immunol 26, 308–322. 10.1038/s41590-024-02059-6.

39. Dimri, G.P., Lee, X., Basile, G., Acosta, M., Scott, G., Roskelley, C., Medrano, E.E., Linskens, M., Rubelj, I., Pereira-Smith, O., and, et al. (1995). A biomarker that identifies senescent human cells in culture and in aging skin in vivo. Proc Natl Acad Sci U S A 92, 9363–9367. 10.1073/pnas.92.20.9363.

40. Zhang, R., Poustovoitov, M.V., Ye, X., Santos, H.A., Chen, W., Daganzo, S.M., Erzberger, J.P., Serebriiskii, I.G., Canutescu, A.A., Dunbrack, R.L., et al. (2005). Formation of MacroH2A-containing senescence-associated heterochromatin foci and senescence driven by ASF1a and HIRA. Dev Cell 8, 19–30.

41. Lojler-Wirth, H., Kalcher, M., and Binder, H. (2015). oposSOM: R-package for high-dimensional portraying of genome-wide expression landscapes on bioconductor. Bioinformatics 31, 3225–3227. 10.1093/bioinformatics/btv342.

42. Kroger, C., Muller, S., Leidner, J., Krober, T., Warnat-Herresthal, S., Spintge, J.B., Zajac, T., Neubauer, A., Frolov, A., Carraro, C., et al. (2024). Unveiling the power of high-dimensional cytometry data with cyCONDOR. Nat Commun 15, 10702. 10.1038/s41467-024-55179-w.

43. Van Gassen, S., Callebaut, B., Van Helden, M.J., Lambrecht, B.N., Demeester, P., Dhaene, T., and Saeys, Y. (2015). FlowSOM: Using self-organizing maps for visualization and interpretation of cytometry data. Cytometry A 87, 636–645. 10.1002/cyto.a.22625.

44. Giles, J.R., Manne, S., Freilich, E., Oldridge, D.A., Baxter, A.E., George, S., Chen, Z., Huang, H., Chilukuri, L., Carberry, M., et al. (2022). Human epigenetic and transcriptional T cell dijerentiation atlas for identifying functional T cell-specific enhancers. Immunity 55, 557–574 e557. 10.1016/j.immuni.2022.02.004.

45. Martos, S.N., Campbell, M.R., Lozoya, O.A., Wang, X., Bennett, B.D., Thompson, I.J.B., Wan, M., Pittman, G.S., and Bell, D.A. (2020). Single-Cell Analyses Identify Dysfunctional CD16+ CD8 T Cells in Smokers. Cell Reports Medicine 1, 100054. 10.1016/j.xcrm.2020.100054.

46. Takeshita, M., Suzuki, K., Kondo, Y., Morita, R., Okuzono, Y., Koga, K., Kassai, Y., Gamo, K., Takiguchi, M., Kurisu, R., et al. (2019). Multi-dimensional analysis identified rheumatoid arthritis-driving pathway in human T cell. Ann Rheum Dis 78, 1346–1356. 10.1136/annrheumdis-2018-214885.

47. Wang, L., Felts, S.J., Van Keulen, V.P., Scheid, A.D., Block, M.S., Markovic, S.N., Pease, L.R., and Zhang, Y. (2018). Integrative Genome-Wide Analysis of Long Noncoding RNAs in Diverse Immune Cell Types of Melanoma Patients. Cancer research 78, 4411–4423. 10.1158/0008-5472.Can-18-0529.

48. Zhang, J.Y., Zhang, Z., Wang, X., Fu, J.L., Yao, J., Jiao, Y., Chen, L., Zhang, H., Wei, J., Jin, L., et al. (2007). PD-1 up-regulation is correlated with HIV-specific memory CD8 T-cell exhaustion in typical progressors but not in long-term nonprogressors. Blood 109, 4671–4678. 10.1182/blood-2006-09-044826.

49. Day, C.L., Kaufmann, D.E., Kiepiela, P., Brown, J.A., Moodley, E.S., Reddy, S., Mackey, E.W., Miller, J.D., Leslie, A.J., DePierres, C., et al. (2006). PD-1 expression on HIV-specific T cells is associated with T-cell exhaustion and disease progression. Nature 443, 350–354. 10.1038/nature05115.

50. Sade-Feldman, M., Yizhak, K., Bjorgaard, S.L., Ray, J.P., de Boer, C.G., Jenkins, R.W., Lieb, D.J., Chen, J.H., Frederick, D.T., Barzily-Rokni, M., et al. (2018). Defining T Cell States Associated with Response to Checkpoint Immunotherapy in Melanoma. Cell 175, 998–1013 e1020. 10.1016/j.cell.2018.10.038.

51. Hazeldine, J., Hampson, P., and Lord, J.M. (2012). Reduced release and binding of perforin at the immunological synapse underlies the age-related decline in natural killer cell cytotoxicity. Aging Cell 11, 751–759. 10.1111/j.1474-9726.2012.00839.x.

52. Wedemeyer, H., He, X.S., Nascimbeni, M., Davis, A.R., Greenberg, H.B., Hoofnagle, J.H., Liang, T.J., Alter, H., and Rehermann, B. (2002). Impaired ejector function of hepatitis C virus-specific CD8 T cells in chronic hepatitis C virus infection. J Immunol 169, 3447–3458. 10.4049/jimmunol.169.6.3447.

53. Kurioka, A., Ussher, J.E., Cosgrove, C., Clough, C., Fergusson, J.R., Smith, K., Kang, Y.H., Walker, L.J., Hansen, T.H., Willberg, C.B., and Klenerman, P. (2015). MAIT cells are licensed through granzyme exchange to kill bacterially sensitized targets. Mucosal Immunol 8, 429–440. 10.1038/mi.2014.81.

54. Johnston, R.J., Comps-Agrar, L., Hackney, J., Yu, X., Huseni, M., Yang, Y., Park, S., Javinal, V., Chiu, H., Irving, B., et al. (2014). The immunoreceptor TIGIT regulates antitumor and antiviral CD8(+) T cell ejector function. Cancer Cell 26, 923–937. 10.1016/j.ccell.2014.10.018.

55. Callan, M.F., Tan, L., Annels, N., Ogg, G.S., Wilson, J.D., O’Callaghan, C.A., Steven, N., McMichael, A.J., and Rickinson, A.B. (1998). Direct visualization of antigen-specific CD8 T cells during the primary immune response to Epstein-Barr virus In vivo. J Exp Med 187, 1395–1402. 10.1084/jem.187.9.1395.

56. Mogilenko, D.A., Shpynov, O., Andhey, P.S., Arthur, L., Swain, A., Esaulova, E., Brioschi, S., Shchukina, I., Kerndl, M., Bambouskova, M., et al. (2021). Comprehensive Profiling of an Aging Immune System Reveals Clonal GZMK(+) CD8(+) T Cells as Conserved Hallmark of Inflammaging. Immunity 54, 99–115 e112. 10.1016/j.immuni.2020.11.005.

57. Majewska, J., Agrawal, A., Mayo, A., Roitman, L., Chatterjee, R., Sekeresova Kralova, J., Landsberger, T., Katzenelenbogen, Y., Meir-Salame, T., Hagai, E., et al. (2024). p16-dependent increase of PD-L1 stability regulates immunosurveillance of senescent cells. Nature cell biology 26, 1336–1345. 10.1038/s41556-024-01465-0.

58. Duraiswamy, J., Ibegbu, C.C., Masopust, D., Miller, J.D., Araki, K., Doho, G.H., Tata, P., Gupta, S., Zilliox, M.J., Nakaya, H.I., et al. (2011). Phenotype, function, and gene expression profiles of programmed death-1(hi) CD8 T cells in healthy human adults. J Immunol 186, 4200–4212. 10.4049/jimmunol.1001783.

59. Tsang, M., Gantchev, J., Ghazawi, F.M., and Litvinov, I.V. (2017). Protocol for adhesion and immunostaining of lymphocytes and other non-adherent cells in culture. Biotechniques 63, 230– 233. 10.2144/000114610.

60. Patel, P.L., and Herbig, U. (2017). Detection of Dysfunctional Telomeres in Oncogene-Induced Senescence. Methods Mol Biol 1534, 69–78. 10.1007/978-1-4939-6670-7_6.

61. Vasilopoulos, T., and Martinez-Zamudio, R.I. (2024). Transcription factor network dynamics during the commitment to oncogene-induced senescence. Frontiers in Epigenetics and Epigenomics 2.

62. Langmead, B., and Salzberg, S.L. (2012). Fast gapped-read alignment with Bowtie 2. Nat Methods 9, 357–359. 10.1038/nmeth.1923.

63. Ritchie, M.E., Phipson, B., Wu, D., Hu, Y., Law, C.W., Shi, W., and Smyth, G.K. (2015). limma powers dijerential expression analyses for RNA-sequencing and microarray studies. Nucleic acids research 43, e47. 10.1093/nar/gkv007.

